# Elimination of separase inhibition reveals absence of cohesin protection in oocyte metaphase II

**DOI:** 10.1101/2025.02.11.637638

**Authors:** Safia El Jailani, Damien Cladière, Elvira Nikalayevich, Sandra A. Touati, Vera Chesnokova, Shlomo Melmed, Eulalie Buffin, Katja Wassmann

## Abstract

The meiotic segregation pattern to generate haploid gametes is mediated by step-wise cohesion removal by separase, first from chromosome arms in meiosis I, and then from the centromere region in meiosis II. In mammalian oocytes, separase is tightly controlled during the hours-long prometaphase and until chromosome segregation in meiosis I, activated for a short time window, and again inhibited until metaphase II arrest is lifted by fertilization. Centromeric cohesin is protected from cleavage by Sgo2-PP2A in meiosis I. It remained enigmatic how tight control of alternating separase activation and inactivation is achieved during the two divisions in oocytes. It was equally unknown when cohesin protection is put in place and removed. Using structure-function assays in knock-out mouse models we established the contributions of cyclin B1 and securin for separase inhibition during both divisions. When eliminating separase inhibition we found that cohesin is not protected early in meiosis I and at metaphase II arrest. Importantly, in meiosis II, the sole event required for cleavage of centromeric cohesin besides separase activation is prior kinetochore individualization in meiosis I.

## Introduction

Correct chromosome segregation during meiosis is essential to generate gametes with the correct ploidy and thus, healthy offspring. Unlike mitosis, during which sister chromatids are segregated to opposite poles to generate two identical daughter cells, meiosis requires segregation of homologous chromosomes in meiosis I and of sister chromatids in meiosis II. Unless pericentromeric cohesin is maintained until meiosis II, sister chromatids separate randomly in anaphase II. This is due to failed establishment of bipolar, tension-bearing attachments with no physical connection between the two sister chromatids. The protease separase is key to remove cohesion holding sister chromatids together. Separase acts by cleaving the kleisin subunit of the cohesin complex, namely Scc1 in mitosis and Rec8 in both meiotic divisions. In meiosis, cohesin cleavage takes place in a step-wise manner: at chromosome arms to resolve chiasmata (sites of recombination) in meiosis I, and at the centromere region in meiosis II. Thus, centromeric cohesin requires to be protected from separase cleavage in meiosis I, and deprotected for proper execution of meiosis II (Marston & Amon, 2004, Petronczki et al., 2003).

Cohesin cleavage in meiosis is controlled by phosphorylation of Rec8, a pre-requisite for efficient separase cleavage, and by inhibitors directly impinging on separase and keeping separase inactive until satisfaction of the spindle assembly checkpoint (SAC) and anaphase onset (Konecna et al., 2023, Wassmann, 2022). Rec8 phosphorylation at the pericentromere in meiosis I is prevented by localization of the phosphatase PP2A-B56 through Sgo2 recruitment, thus protecting Rec8 from cleavage during the first meiotic division (Gutierrez-Caballero et al., 2012, Marston, 2015). We recently showed that a physical separation of sister kinetochores (sister kinetochore individualization) prior to meiosis I-to-meiosis II transition is key for deprotection and cleavage of centromeric cohesin in meiosis II (Gryaznova et al., 2021). In budding yeast, centromeric cohesin protection is eliminated at the metaphase-to-anaphase transition of meiosis II in an APC/C (Anaphase Promoting Complex/ Cyclosome)-dependent manner (Jonak et al., 2017, Mengoli et al., 2021). It is yet unclear whether or not this also occurs in oocytes, which have to maintain an extended arrest in metaphase II to await fertilization, and whether cohesin deprotection in oocytes requires APC/C activity.

In terms of the cell-cycle, the meiotic divisions constitute a real challenge, as cells have to execute two M-phases without interphase and the associated reset of the cell cycle in-between. Additionally, oocytes await fertilization during a prolonged metaphase II arrest of variable length, before lift of the arrest and anaphase II onset only upon sperm entry (Holt et al., 2013). Oocytes are not renewed from birth of the female until fertilization, placing incredibly temporal strain on female gametes. This leads to the deterioration of cohesin complexes and SAC control, and the generation of aneuploidies with increasing maternal age (Herbert et al., 2015). It is therefore important to understand how separase activity, cell cycle progression and step-wise cohesin removal are coordinated in mammalian oocyte meiosis.

Separase has to be tightly controlled until anaphase I onset, transiently activated to allow removal of arm cohesin, and re-inhibited for entry into meiosis II. Again, at entry into meiosis II, re-inhibition of separase must be very tight but at the same time readily reversible when fertilization occurs (Konecna et al., 2023, Wassmann, 2022). In both mitosis and meiosis I prophases, separase is excluded from the nucleus due to a nuclear export signal (Hellmuth et al., 2018, Sun et al., 2006). Thus, in mitosis, separase inhibition becomes important once nuclear envelope breakdown occurs. Importantly, in oocytes, separase is not physically separated from its substrate at the transition from meiosis I into meiosis II, because no nucleus is reformed between the two meiotic divisions. It is currently unclear how separase inhibition is assured when oocytes progress from meiosis I into meiosis II.

To determine when cohesin protection and deprotection is present during meiosis, it is necessary to establish how separase activity is controlled, to generate conditions where active separase can cleave cohesin that is not protected, at the different meiotic cell cycle stages. Cyclin B1 and securin, which both inhibit vertebrate separase in mitosis and meiosis, are ubiquitinated by the APC/C and thus targeted for degradation prior to anaphase onset (Kamenz & Hauf, 2017, Yu et al., 2023). Of note, securin, which was discovered as pituitary tumor transforming gene (PTTG) (Pei & Melmed, 1997), is both a separase inhibitor and a chaperone for separase, and therefore contributes to full separase activity (Holland & Taylor, 2008). For separase inhibition, securin acts as a pseudosubstrate and blocks separase by binding to its substrate pocket (Lin et al., 2016, Waizenegger et al., 2002, Yu et al., 2021). Cyclin B1, together with Cdk1, inhibits separase through the phosphorylation of a conserved residue, creating a pocket for the inhibitory binding of cyclin B1-Cdk1 to separase (Gorr et al., 2005, Holland & Taylor, 2006, Stemmann et al., 2001, Yu et al., 2021). Separase inhibition by securin or cyclin B1 is mutually exclusive. Initially, it was proposed that cyclin B1-dependent inhibition of separase occurs concomitantly with securin (Stemmann et al., 2001). Alternatively, cyclin B1 inhibition of separase may become important once securin is degraded, because complete cyclin B1 degradation is temporally delayed compared to securin (Afonso et al., 2019, Collin et al., 2013, Shindo et al., 2012, Wolf et al., 2006, Yu et al., 2023). Consistent with a hand-over from securin to cyclin B1-Cdk1-dependent separase control (Yu et al., 2023), cyclin B1-Cdk1 activity itself is also inhibited when bound to separase, which may thus contribute to down-regulation of Cdk1 activity at exit from mitosis and at the transition from meiosis I into meiosis II (Gorr et al., 2006). Nevertheless, in the absence of securin, cyclin B1-Cdk1 may phosphorylate and inhibit securin-free separase also in mitotic prometaphase, explaining why loss of securin -either in tissue culture cells or in a mouse model-is not lethal (Jallepalli et al., 2001, Mei et al., 2001, Pfleghaar et al., 2005, Wang et al., 2003, Wang et al., 2001).

Both inhibitors control separase activity in oocyte meiosis, and degradation of both is required for metaphase-to-anaphase transition in meiosis I (Herbert et al., 2003, Terret et al., 2003). Securin was proposed to function as the main separase inhibitor in oocyte meiosis II (Nabti et al., 2008), at odds with the fact that mice null for securin are viable and fertile (Mei et al., 2001, Wang et al., 2001). Together, the relative contributions and potential redundancies of separase inhibition by either securin or cyclin B1 in meiosis I and meiosis II are still unknown.

A third inhibitor of separase, namely Sgo2-Mad2, is present in somatic cells arrested in mitosis due to SAC activation (Hellmuth et al., 2020). Whether this inhibitor has a physiological role in cycling cells or in meiosis to keep separase in check, is currently unknown. Sgo2 occupies multiple roles in oocyte meiosis, beyond centromeric cohesin protection. Complete loss of Sgo2 leads to delayed anaphase I onset, due to its function in silencing the spindle checkpoint. Segregation occurs only after APC/C activation in meiosis I, and there are no indications for precociously activated separase without Sgo2 (Marston, 2015, Rattani et al., 2013).

It has been proposed that pericentromeric cohesin protection is maintained until metaphase II to prevent separase from cleaving this fraction of cohesin holding sister chromatids together, as separase may become reactivated upon degradation of cyclin B1 and securin at meiosis I exit. However, this hypothesis is not consistent with most Sgo2 supposedly required for PP2A recruitment and cohesin protection being displaced from pericentromeric chromatin at meiosis I exit before being recruited there again at high levels as oocytes progress into metaphase II (Gryaznova et al., 2021, Mengoli et al., 2021). Several models have been proposed for deprotection of centromeric cohesin in metaphase II: Bipolar tension applied by the spindle on sister kinetochores in meiosis II was thought to lead to the deprotection of centromeric cohesin, e.g. at a time when oocytes have been able to re-accumulate separase inhibitor(s) (Gomez et al., 2007, Lee et al., 2008), however this “deprotection by tension model” was not confirmed (Gryaznova et al., 2021, Mengoli et al., 2021). In budding yeast, APC/C activation leading to the degradation of Sgo1 and Mps1 was shown to mediate deprotection, but whether this also applies to oocytes was unknown (Jonak et al., 2017, Mengoli et al., 2021). We previously reported a role for Set/I2PP2A in promoting cohesin removal in meiosis II (Chambon et al., 2013); however, our recent, unpublished results show that Set/I2PP2A does so independently of cohesin protection (Keating et al., in preparation). More recently we found that sister kinetochore individualization by separase in anaphase I is key for cleavage of centromeric cohesin in the subsequent meiosis II division (Gryaznova et al., 2021), but it is unclear whether kinetochore individualization is required to remove protection still present, or simply allow access to pericentromeric Rec8. Thus, it is unknown when pericentromeric cohesin protection is present, and when it is removed after metaphase-to-anaphase transition in oocyte meiosis I.

Here, we set out to determine the contributions of the two main separase inhibitors, cyclin B1-Cdk1 and securin, for maintaining separase control during the first and second meiotic division in mammalian oocytes, and to understand when exactly centromeric cohesin is protected from cleavage. Separase inhibition in oocytes is dose-dependent (Chiang et al., 2011) and it is thus difficult to obtain conclusive results with transient knock-down approaches and exogenous protein expression alone. Using knock-out mouse strains and rescue experiments, we found that either cyclin B1-Cdk1 or securin are sufficient to maintain separase in check up until metaphase in both meiosis I and II. However, at the transition from meiosis I into meiosis II, cyclin B1-Cdk1 rather than securin is essential to re-inhibit separase. Strikingly, loss of both inhibitory mechanisms in either meiotic division leads to immediate separase activation and cleavage of cohesin. Importantly, without securin and cyclin B1-dependent inhibition of separase we determined when centromeric cohesin protection is in place during oocyte meiotic divisions. Unexpectedly, we found that centromeric cohesin protection is not yet set-up after nuclear envelope breakdown in early prometaphase I. Additionally, protection is not maintained during the meiosis I into meiosis II transition. Surprisingly, protection is also absent during the metaphase II arrest, when oocytes await fertilization. Hence, mechanisms proposed for deprotection of centromeric cohesin in metaphase II promote cleavage independently of cohesin protection. Our results depict critical moments during meiotic cell cycle progression prone to missegregation events and the generation of aneuploidies in the absence of tight separase control in oocytes.

## Results

### Loss of cyclin B1-mediated separase inhibition does not disturb bivalent segregation

We have shown previously that exogenously expressed separase carrying a nonphosphorylatable amino-acid substitution of the conserved residue targeted by cyclin B1-Cdk1 (separase S1121A) is not inhibited by cyclin B1 anymore during oocyte meiosis I (Touati et al., 2012). However, it was unknown whether cyclin B1-dependent inhibition of separase plays an important role in a physiological context (i.e., in presence of securin) up to metaphase I and beyond.

To clarify this issue in an undisputable manner we performed structure-function assays through complementation, using separase knock-out mouse oocytes. Mature oocytes can be harvested before resumption of meiosis I, in prophase I (also called GV stage). They can be injected with mRNAs to express proteins of our choice, and induced to undergo meiosis I in a synchronized manner in culture. After GVBD (Germinal Vesicle Breakdown, corresponds to nuclear envelope breakdown) oocytes undergo prometaphase and reach metaphase around 6 hours after GVBD. Metaphase-to-anaphase transition of meiosis I with polar body (PB) extrusion takes place around 8 hours after GVBD. This is followed by exit from meiosis I and progression into meiosis II. Oocytes reach metaphase II approximately 12 hours after GVBD and remain arrested for up to 12 hours (mouse), until fertilization occurs. Fertilization induces anaphase II onset and exit from meiosis II (**Figure 1A**).

**Figure 1:**
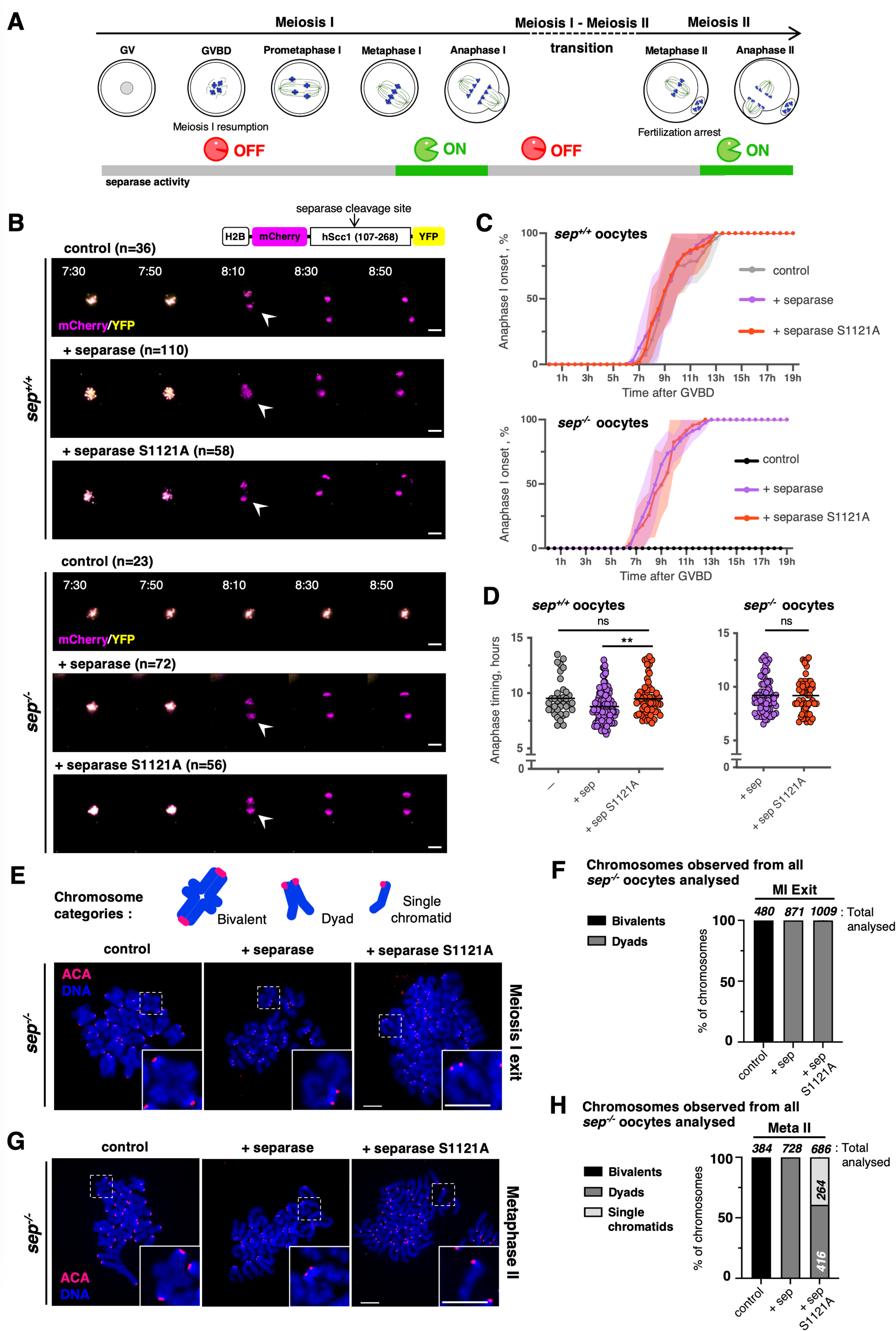
Cyclin B1-dependent inhibition of separase becomes essential only after meiosis I exit. **A)** Scheme of mouse oocyte meiosis I and II. Chromosomes are in blue, spindles in green, and the nucleus (Germinal Vesicle, GV) in grey. Separase activity at the different cell cycle stages is indicated below. **B)** On top right, schematic design of the Scc1 cleavage sensor construct used. Below, overlays of the YFP and mCherry channels of time lapse microscopy acquisitions of wild type (*sep^+/+^*) and separase knock-out (*sep^−/−^*) mouse oocytes expressing the cleavage sensor. Time after GVBD is shown in hours:minutes (anaphase I onset is indicated with an arrow head). Oocytes have been co-injected with mRNAs encoding for separase or separase S1121A, where indicated. Cleavage of the sensor is visible by the disappearance of the YFP signal from the chromosomes, whereas the mCherry signal remains localized to chromosomes. n is the number of oocytes analysed. Scale bar (white) represents 20 μm. **(related to Figure EV1)**. **C)** Monitoring anaphase I onset of the selected time frames shown in (B). Hours after germinal vesicle breakdown (GVBD) are indicated. For each graph, error bars are ± SD. **D)** Statistical analysis of anaphase timing of *sep^+/+^* and *sep^−/−^* oocytes from (B). For each graph mean is shown, ns indicate there is no significant difference, and asterisks indicate significant difference (** P<0.01) according to Mann-Whitney U-test. **E)** Top: representative schemes of chromosome categories observed, classified as “bivalent” (paired homologous chromosomes), “dyad” (paired sister chromatid), or “single chromatid”. Bottom: *sep^−/−^* oocytes expressing separase or separase S1121A were fixed for chromosome spreads around 8h after GVBD (Meiosis I exit). Oocytes were fixed after visual selection of oocytes extruding the polar body (PB), collection of *sep^−/−^* oocyte was time-matched. Centromeres/kinetochores were stained with ACA (red) and chromosomes with DAPI (blue). A representative spread and magnification of one chromosome (white dashed line squares, insert at the bottom right corner) are shown for each condition. Total number of chromosomes analysed is indicated in (F). Scale bars (white) represent 10 μm. **F)** Frequency of chromosome categories observed at exit from meiosis I (MI Exit), quantified from chromosome spreads shown in (E). Where indicated, *sep^−/−^* oocytes express separase or separase S1121A. The total number of chromosomes quantified for each condition is indicated. sep: separase. **G)** *sep^−/−^* mouse oocytes expressing separase or separase S1121A were fixed for chromosome spreads around 20h after GVBD (Metaphase II). *sep^−/−^* oocytes were collected at the same time. Centromeres/kinetochores were stained with ACA (red) and chromosomes with DAPI (blue). A representative spread and magnification of one chromosome (white dashed line squares, insert at the bottom right corner) are shown for each condition. Total number of chromosomes analysed is indicated in (H). Scale bars (white) represent 10 μm. sep: separase. **H)** Frequency of chromosome categories observed at metaphase II arrest (MetaII), quantified from chromosome spreads in (G). Where indicated, *sep^−/−^* oocytes express separase or separase S1121A. The total number of chromosomes quantified for each condition, and of each category, is indicated.

A mouse strain allowing the conditional oocyte-specific invalidation of separase was obtained through the Cre-LoxP system under the control of the oocyte specific *Zona pellucida 3* (Zp3) promoter. This strain has been previously characterized (Kudo et al., 2006). For simplicity, oocytes of *separase^LoxP/LoxP^ Zp3 Cre^+^* mice are designated as *sep^−/−^*, and controls (*separase^LoxP/LoxP^*) as *sep^+/+^*. Oocytes deleted for endogenous separase were thus used to address whether expression of the separase S1121A mutant caused inappropriate separase activation before metaphase I or failure to re-inhibit separase after anaphase I.

Control (*sep^+/+^*) and *sep^−/−^* oocytes were injected with mRNA coding either for wild type separase or separase S1121A. To follow separase activity by live imaging we additionally expressed a separase activity sensor (Nikalayevich et al., 2018), based on a similar sensor initially described (Shindo et al., 2012). This cleavage biosensor contains a separase cleavage site and is artificially localized to chromosomes. Upon cleavage of the sensor, co-localization of the two fluorochromes of the sensor is lost, leading to a change of color that allows us to determine separase activation as well as anaphase onset timing, because the sensor allows us to follow chromosome movements as well **(movie EV1)**. Whereas no cleavage of the sensor was observed in oocytes devoid of functional separase, cleavage of the sensor took place with similar timing in *sep^−/−^* oocytes expressing wild type or S1121A mutant separase **(Figure 1B-D, Figure EV1)**. Of note, a small but still significant delay in anaphase I onset was observed when separase was expressed in addition to endogenous separase in wild type but not in *sep^−/−^* oocytes **(Figure 1D)**. The reasons for this are unclear, but may be related to mutual inhibition of separase and cyclin B1-Cdk1 activity, reducing endogenous Cdk1 activity required for progression through meiosis I, when wild type separase is expressed additionally to endogenous separase (Gorr et al., 2005, Gorr et al., 2006, Shindo et al., 2012). Crucially, chromosome spreads performed after metaphase to anaphase transition of meiosis I confirmed that bivalent chromosomes had been segregated into dyads under the rescue conditions, no matter whether wild type or S1121A separase was expressed **(Figure 1E and F)**, and similar to what we had observed before. Precocious sister chromatid segregation was not observed. These results indicate that cyclin B1-dependent inhibition of separase alone is not essential to correctly inhibit separase, at least until chromosome segregation occurs in meiosis I.

### Cyclin B1-Cdk1 becomes essential to inhibit separase after anaphase I

We asked whether re-inhibition of separase after meiosis I, occurred correctly in *sep^−/−^* oocytes expressing S1121A mutant separase. This is an important question, because it has been observed that Sgo2, which is required for protection of centromeric cohesin, disappears from the centromere region in late anaphase I, before strongly reaccumulating there again in meiosis II (Gryaznova et al., 2021, Mengoli et al., 2021). Furthermore, sister kinetochore individualization, a pre-requisite for centromeric cohesin cleavage in meiosis II, also occurs already in late anaphase I (Gryaznova et al., 2021). Therefore, re-inhibition of separase is most likely important to avoid cleavage of potentially unprotected cohesin and as a consequence, precocious sister chromatid segregation at the transition from meiosis I into meiosis II. To answer this question, we repeated the previous rescue experiment, but performed chromosome spreads at a later timepoint, when oocytes had reached metaphase II and were awaiting fertilization. Crucially, now we observed a high fraction of single sister chromatids in *sep^−/−^* oocytes rescued with S1121A separase but not with wild type separase, indicating that during the transition from meiosis I to meiosis II, separase remained active when cyclin B1-dependent inhibition was absent **(Figure 1G and H**). These data also indicate that centromeric cohesin protection was absent, leading to cleavage of cohesin at the centromere region. We conclude that cyclin B1-dependent inhibition of separase is essential once anaphase I has taken place and oocytes progress into meiosis II.

### Securin is not essential in oocyte meiosis I

The nature of the inhibitory mechanism used by securin is not compatible with the generation of separase mutants that would only interfere with securin inhibition (Yu et al., 2023). For this reason, we decided to use oocytes derived from a complete securin knock-out (*Pttg^−/−^* mice (Wang et al., 2001), for clarity we call oocytes devoid of securin, *securin^−/−^* oocytes). Mice devoid of securin are viable, exhibit features of senescence, and demonstrate some signs of female subfertility (Chesnokova & Melmed, 2010, Mei et al., 2001, Wang et al., 2001). Oocytes from these complete knock-out mice resumed meiosis without delay. Separase activation and chromosome segregation occurred on time and without any noticeable issues, indicating that securin is not essential in meiosis I **(Figure 2A and B, Figure EV2)**. A small delay in anaphase timing was observed in s*ecurin^−/−^* oocytes; however, it was not statistically significant **(Figure 2C)**, indicating that the chaperone activity of securin is not important for full separase activity in oocyte meiosis (Holland & Taylor, 2008). Hardly any precocious sister chromatid segregation was observed when *securin^−/−^* oocytes reached metaphase II arrest, indicating that securin is also not essential during the transition from meiosis I into meiosis II **(Figure 2D and E)**.

**Figure 2.**
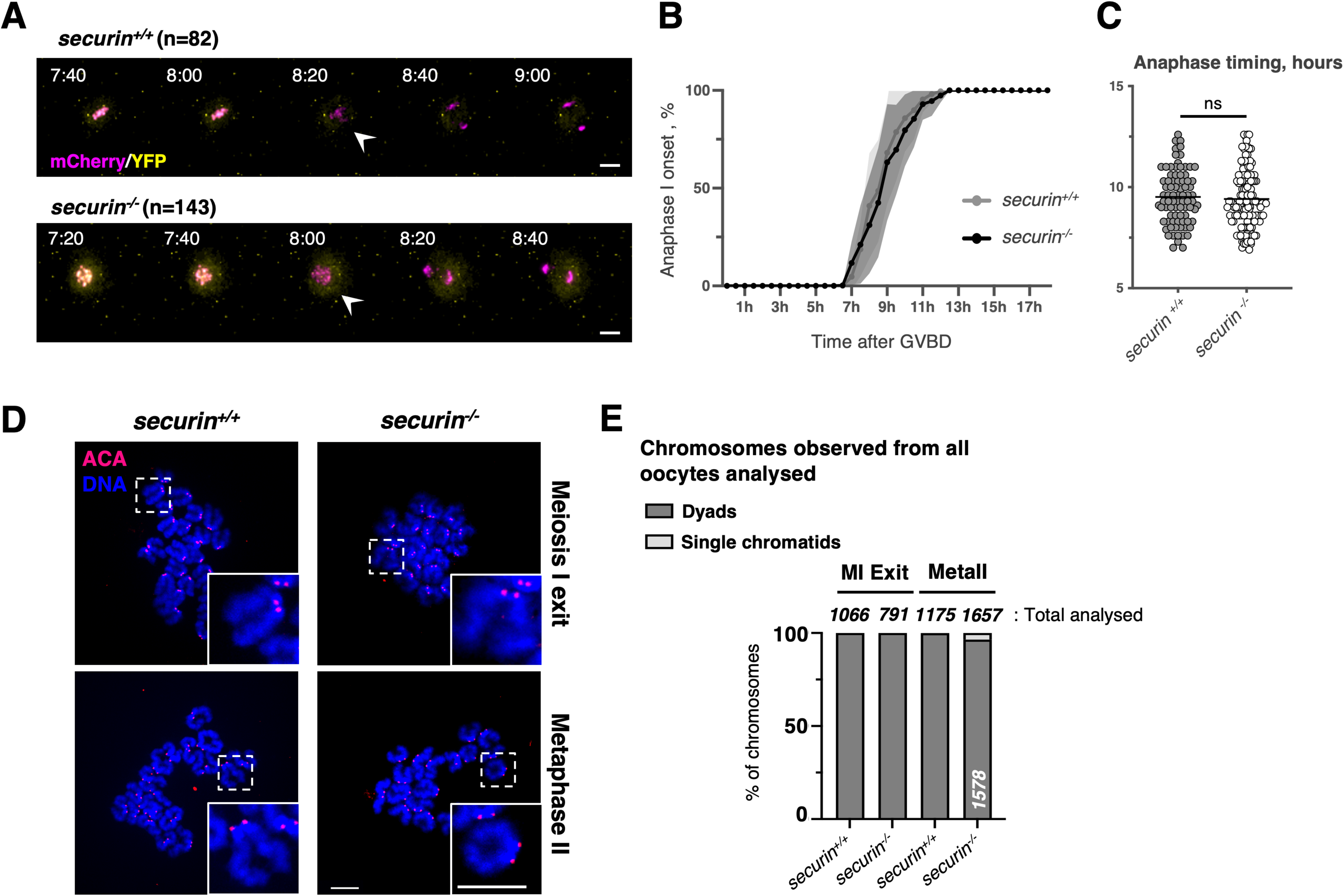
Securin is not essential for separase inhibition in meiosis I. **A)** Overlays of the YFP and mCherry channels of time lapse microscopy acquisition such as described in Fig 1A, of wild type (*securin^+/+^*) and securin knock-out (*securin^−/−^*) oocytes expressing the cleavage sensor. Time after GVBD is shown in hours:minutes (anaphase I onset is indicated with an arrow head). n is the number of oocytes analysed. Scale bar (white) represents 20 μm. **(related to Figure EV2)** **B)** Monitoring anaphase I onset in the corresponding conditions of the selected time frames shown in (A). Hours after GVBD are indicated, and error bars are ± SD. **C)** Statistical analysis of anaphase timing of *securin^+/+^* and *securin^−/−^* oocytes from (A). For each graph mean is shown, and ns indicate there is no significant difference according to Mann-Whitney U-test. **D)** *securin^+/+^* and *securin^−/−^* oocytes were fixed for chromosome spreads around 8h after GVBD (Meiosis I Exit) and 20h after GVBD (Metaphase II). Centromeres/kinetochores were stained with ACA (red) and chromosomes with DAPI (blue). A representative spread and magnification of one chromosome (white dashed line squares, insert at the bottom right corner) are shown for each condition. Total number of chromosomes analysed is indicated in (E). Scale bars (white) represents 10 μm. **E)** Frequency of chromosome categories observed at exit from meiosis I (8 hours after GVBD) and at metaphase II arrest (20 hours after GVBD), quantified from chromosome spreads in (D). The total number of chromosomes quantified for each condition, and of each category, is indicated.

### “Separase-out-of-control” phenotype in prometaphase I

Are securin and cyclin B1 together the main separase inhibitors in meiosis? To clarify this issue, we created *separase^LoxP/LoxP^ Zp3 Cre^+^ Pttg^−/−^* mice. Oocytes are thus devoid of separase and securin (s*ep^−/−^ securin^−/−^* oocytes) and can be used for rescue experiments with separase S1121A. In these oocytes, separase is free of cyclin B1 and securin inhibition.

First, we asked how oocytes progress through meiosis I in absence of both separase inhibitors. In s*ep^−/−^ securin^−/−^* oocytes rescued with wild type separase, separase activation took place on time and anaphase I was visible. Strikingly, when separase S1121A was expressed, separase became activated immediately after resumption of meiosis I, shortly after germinal vesicle breakdown (GVBD), which corresponds to nuclear envelope breakdown **(Figure 3A)**. s*ep^−/−^ securin^−/−^* oocytes rescued with separase S1112A and followed by live imaging showed chaotic chromosome segregation taking place, and cleavage of the separase sensor started shortly after GVBD, a phenotype we coined “separase-out-of-control” **(Figure 3A-C** (compare also to **Figure 1B), Figure EV3, movie EV2,** for comparison of stills of entire movies see **Appendix Figure S1)**. We conclude that separase must be inhibited by either cyclin B1 or securin in oocyte meiosis I, and that either inhibitor is sufficient, but when both are absent, separase immediately cleaves cohesin.

**Figure 3.**
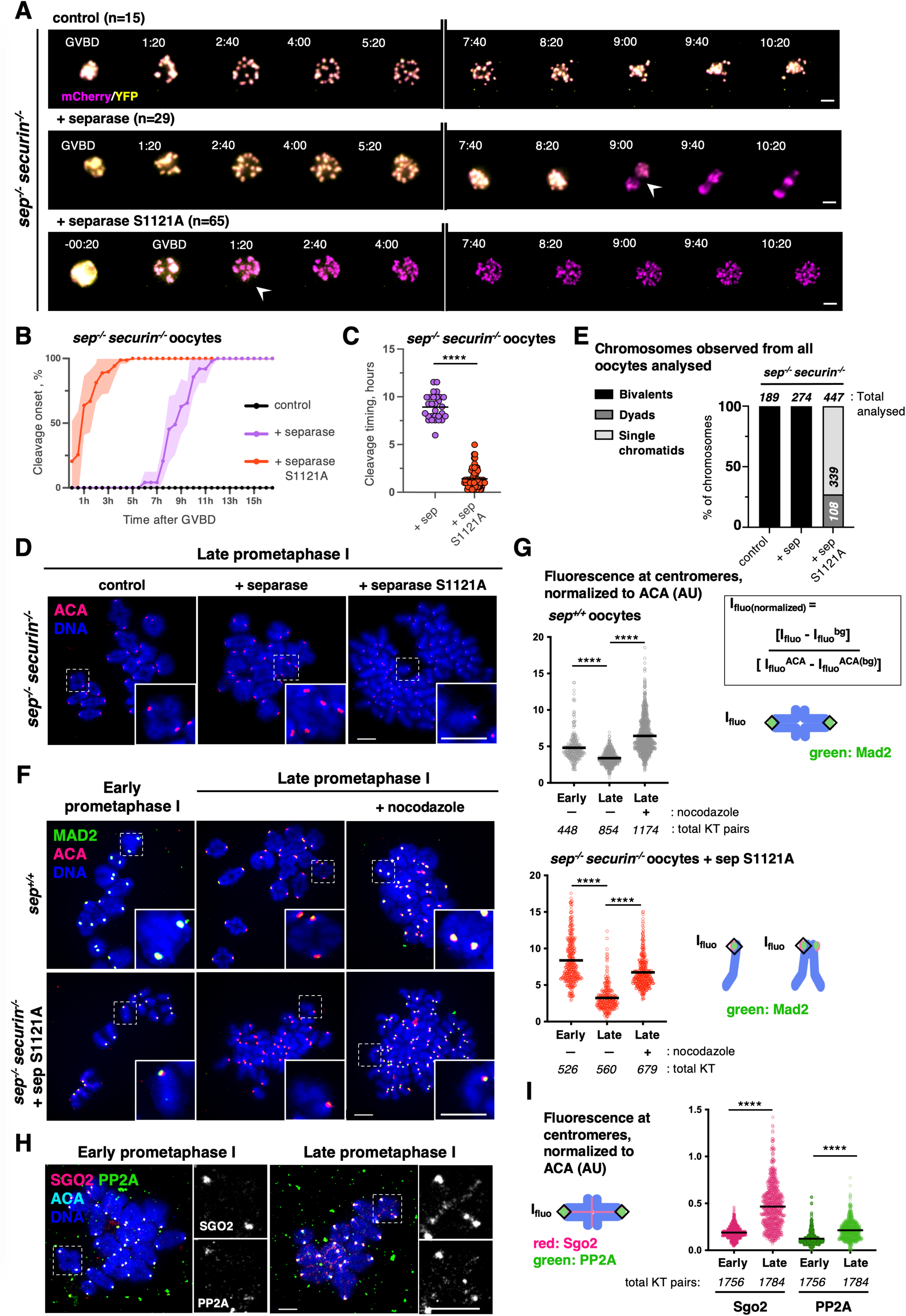
Complete loss of separase control reveals absence of cohesin protection at meiosis resumption. **A)** Overlays of the YFP and mCherry channels of time lapse microscopy acquisition such as described for Fig 1A, of *sep^−/−^securin^−/−^* oocytes expressing the cleavage sensor, from GVBD onwards. Time after GVBD is shown in hours:minutes (cleavage onset is indicated with an arrow head). Oocytes have been co-injected with mRNAs encoding for separase or separase S1121A, where indicated. n is the number of oocytes analysed. Scale bar (white) represents 20 μm. **(Related to Figure EV3, movie EV2 and Appendix Fig. S1)** **B)** Monitoring of the switch of colour indicative of cleavage onset, in the corresponding conditions of the selected time frames shown in (A). Hours after GVBD are indicated, and error bars are ± SD. **C)** Statistical analysis of cleavage timing in *sep^−/−^securin^−/−^* oocytes from (A). For each graph, mean is shown, and asterisks indicate significant difference (**** P<0.0001) according to Mann-Whitney U-test. **D)** *sep^−/−^securin^−/−^* oocytes expressing separase or separase S1121A were fixed for chromosome spreads around 5h after GVBD (Late prometaphase I). Centromeres/kinetochores were stained with ACA (red) and chromosomes with DAPI (blue). A representative spread and magnification of one chromosome (white dashed line squares, insert at the bottom right corner) are shown for each condition. Total number of chromosomes analysed is indicated in (E). Scale bars (white) represents 10 μm. **E)** Frequency of chromosome categories observed around 5h after GVBD (Late prometaphase I), quantified from chromosome spreads shown in (D). The total number of chromosomes quantified for each condition, and of each category, is indicated. **F)** Not injected *sep^+/+^* (top) or *sep^−/−^ securin^−/−^* (bottom) oocytes injected in GV with separase S1121A were fixed for chromosome spreads around 3h after GVBD (Early prometaphase I) and around 6h after GVBD (Metaphase I). Oocytes were treated with nocodazole at GVBD, where indicated. Chromosome spreads were stained for endogenous Mad2 (green), centromeres/kinetochores were stained with ACA (red) and chromosomes with DAPI (blue). A representative spread and magnification of one chromosome (white dashed line squares, insert at the bottom right corner) are shown for each condition. Scale bars (white) represents 10 μm. **G)** Total Mad2 fluorescence intensities at centromeres/kinetochores were quantified from chromosome spreads in (F) in not injected *sep^+/+^* (top) and *sep^−/−^ securin^−/−^* (bottom) oocytes expressing separase S1121A. Where indicated, oocytes were treated with nocodazole at GVBD. The signals measured (Ifluo) were corrected to background (bg) and normalized to ACA signals. On the graph, mean is shown, and asterisks indicate significant difference (**** P<0.0001) according to Mann-Whitney U-test. The number of kinetochore pairs (KT pairs) and kinetochore (KT) used for quantification is indicated. On the right, scheme of Mad2 (green) signal measurement. AU, arbitrary units. **H)** Wild type oocytes were fixed for chromosome spreads at 1h30 after GVBD (early prometaphase I) and 4h after GVBD (late prometaphase I). Chromosome spreads were stained for endogenous Sgo2 (red), endogenous PP2A (green), centromeres/kinetochores were stained with ACA (cyan) and chromosomes with DAPI (blue). Representative spreads and magnifications of one chromosome (white dashed line squares) are shown for each condition, for Sgo2 (top) and PP2A (bottom) total staining. Scale bars (white) represents 10 μm. **I)** Total Sgo2 (red) and total PP2A (green) fluorescence intensities at centromeres/kinetochores were quantified from chromosome spreads in (H). The signals measured (Ifluo) were corrected to background (bg) and normalized to ACA signals. On the graph, mean is shown, and asterisks indicate significant difference (**** P<0.0001) according to Mann-Whitney U-test. The number of kinetochore pairs (KT pairs) used for quantification is indicated. On the right, scheme of Sgo2 (red) and PP2A (green) signal measurements. AU, arbitrary units.

### Absence of separase control reveals absence of cohesin protection in early prometaphase I

Importantly, chromosome spreads in late prometaphase I revealed that the majority of bivalent chromosomes had segregated into sister chromatids when *sep^−/−^ securin^−/−^* oocytes expressed separase S1121A. This was not the case when they expressed wild type separase, which can still be inhibited by cyclin B1 **(Figure 3D and E)**. Presence of active separase allows us to conclude that centromeric cohesin is not protected at meiosis I resumption. Because cohesin gets cleaved so early, no real anaphase movement was observed **(Figure 3A, movie EV2)**.

The absence of anaphase I movements in “separase-out-of control” oocytes suggested that under these conditions, separase is activated and able to cleave all cohesin at a time when the APC/C is still inactive. To confirm that chromosomes and sister chromatids indeed separate in absence of APC/C activation, we asked whether the SAC, which inhibits the APC/C, is active at the time of separase cleavage. In oocytes, it has been shown that the SAC is activated from GVBD until around 4 hours after GVBD because of missing stable end-on attachments. The SAC gets progressively shut off as oocytes progress into metaphase I. This is revealed by staining for the SAC protein Mad2, which gets recruited to unattached kinetochores (Kitajima et al., 2011, Lane et al., 2012, Wassmann et al., 2003). To establish whether separase cleaves centromeric and arm cohesin even though the APC/C is still inactive, we stained chromosome spreads for Mad2. **Figure 3F and G** shows that in both controls and “separase-out-of-control” oocytes, Mad2 is present at kinetochores in prometaphase I (3 hours after GVBD). Strikingly, single sister chromatids in *sep^−/−^ securin^−/−^* oocytes expressing separase S1121A showed a strong Mad2 signal at kinetochores. Mad2 staining was reduced in metaphase I, even in *sep^−/−^ securin^−/−^* oocytes expressing separase S1121A, indicating that the SAC, which is only activated transiently in oocytes, had been turned off also in the presence of single sister chromatids. Treating “separase-out-of-control” oocytes with nocodazole at resumption of meiosis and at a concentration that prevents chromosome segregation in control oocytes, led to strong Mad2 recruitment. However, nocodazole treatment did not prevent cohesin cleavage and sister chromatid separation, further indicating that it took place independently of APC/C activation (**Figure 3F and G)**. We conclude that loss of securin and cyclin B1-mediated inhibition of separase leads to cleavage of all cohesin in an APC/C-independent manner upon meiosis I resumption.

Our results indicate that centromeric cohesin protection is absent when oocytes resume meiosis I. We hypothesized that separase activation happens so early in s*ep^−/−^ securin^−/−^* oocytes rescued with separase S1121A, that cohesin protection is not yet in place, leading to the observed phenotype. To further address this, we examined whether the cohesin protectors Sgo2 and PP2A are localized to the centromere region at all, early in meiosis I. Indeed, when checking for localization of Sgo2 and PP2A to the centromere region in early prometaphase I of wild type oocytes, we found that their recruitment to centromeres increases with time and is significantly lower in early than in late prometaphase I **(Figure 3H and I)**. Centromeric cohesin protection is thus being gradually set up as oocytes progress through prometaphase of meiosis I. We conclude that the presence of active separase at meiosis resumption leads to the observed cleavage of all cohesin, on arms and at the centromere region, because centromeric cohesin is not yet protected at that time.

### Cyclin B1 is not the major inhibitor of separase in meiosis II

In a previous study, using injection experiments with blocking antibodies and transient knock-down approaches, securin and not cyclin B1-Cdk1 was proposed as the major separase inhibitor in oocyte meiosis II (Nabti et al., 2008). Using *sep^−/−^* oocytes, we set out to determine whether cyclin B1 alone is indeed not required for separase inhibition in meiosis II. As we have shown previously, *sep^−/−^* oocytes cannot separate chromosomes in meiosis I, however, as chromosomes are correctly attached, the SAC is satisfied and the APC/C is activated (Kudo et al., 2006) These oocytes progress through meiosis I into meiosis II, with the degradation and re-accumulation of APC/C substrates. Hence, *sep^−/−^* oocytes that have reached meiosis II can be injected with mRNA coding for wild type separase or S1121A separase to ask whether chromosome separation is observed, indicating separase activation (Gryaznova et al., 2021). Thus, this experimental tool allows us to address whether separase inhibition by cyclin B1 becomes important during the prolonged metaphase II arrest that oocytes are subjected to when they are awaiting fertilization **(Figure 4A)**.

**Figure 4.**
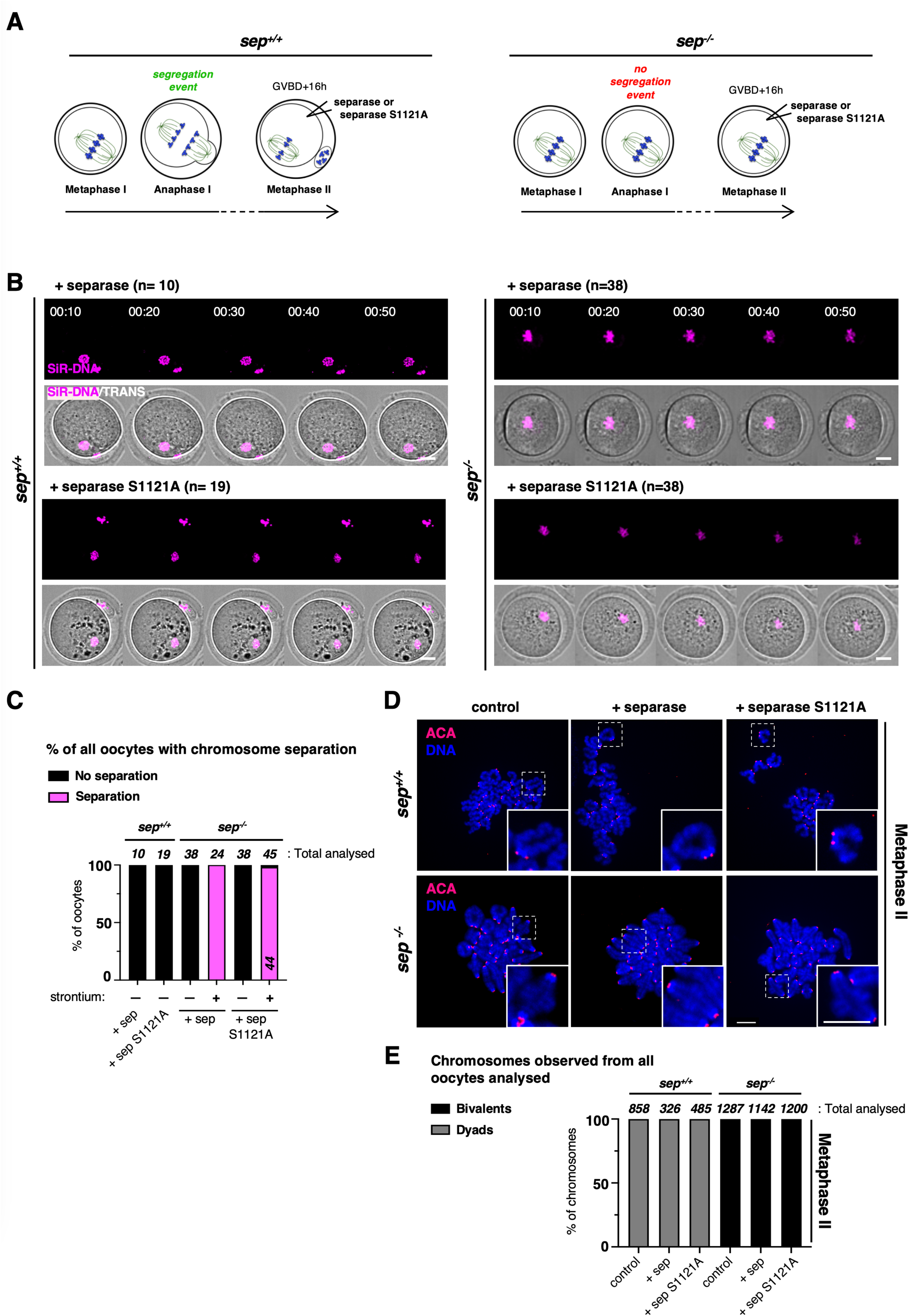
Inhibition of separase by cyclin B1 or securin alone is not essential in meiosis II. **A)** Illustrative scheme of the experimental setup used. Oocytes of the indicated genotype were matured *in vitro* until metaphase II. *sep^+/+^* correspond to *separase^LoxP/LoxP^*. Of note, *sep^−/−^* oocytes progress into metaphase II, even though no chromosome segregation occurs in meiosis I (Kudo, 2006). Once arrested at metaphase II, oocytes were injected with mRNAs encoding for separase or separase S1121A. Post injections, oocytes were either subjected to live imaging to track chromosome segregation (B), or chromosome spreads were performed to verify cohesin cleavage (D). **B)** Live imaging of *sep^+/+^* (left) or *sep^−/−^* (right) oocytes undergoing meiosis II (from metaphase II onwards) and expressing separase or separase S1121A. For all conditions, oocytes were injected around 16h after GVBD (arrested in metaphase II) and were subjected to live imaging around 18h after GVBD. On top, the magenta channel to visualize chromosomes, and below overlay with TRANS channel image. Time after start of the movie is shown in hours:minutes. Prior to acquisition, oocytes were preincubated in culture media containing SiR-DNA to visualize chromosomes. Efficient expression of the separase constructs was verified by inducing anaphase II chemically with strontium (**Related to Figure EV4A**). n is the number of oocytes analysed. Scale bar (white) represents 20 μm. **C)** Frequency of DNA separation observed in metaphase II-arrested oocytes, indicating separase activation, in (B). The total number of oocytes analysed for each condition, and of each category, is indicated. sep: separase. (**Related to Figure EV4A**) **D)** *sep^+/+^* (top) or *sep^−/−^* (bottom) oocytes injected around 16h after GVBD (arrested in metaphase II) with the indicated constructs, were fixed for chromosome spreads around 22h after GVBD (Metaphase II). Centromeres/kinetochores were stained with ACA (red) and chromosomes with DAPI (blue). A representative spread and magnification of one chromosome (white dashed line squares, insert at the bottom right corner) are shown for each condition. Total number of chromosomes analysed is indicated in (E). Scale bars (white) represents 10 μm. **E)** Frequency of chromosome categories observed at metaphase II arrest (22h after GVBD), quantified from chromosome spreads shown in (D). The total number of chromosomes quantified for each condition, is indicated. sep: separase.

As expected, injection of wild type separase into control or *sep^−/−^* oocytes did not lead to chromosome or sister chromatid separation during metaphase II arrest, because separase remained correctly inhibited **(Figure 4B-E)**. Importantly, the same was true when separase S1121A was expressed, showing that phosphorylation of S1121 by cyclin B1-Cdk1 is not essential, or not the sole mechanism, required to keep separase under control during the metaphase II arrest **(Figure 4B-E)**. Thus, cyclin B1 is the sole mechanism inhibiting separase at the transition from meiosis I into meiosis II (**Figure 1G and H**), but once oocytes have reached metaphase II, loss of cyclin B1 inhibition alone was not enough to activate separase. Mimicking fertilization through strontium treatment induced anaphase II onset in control and *sep^−/−^* oocytes **(Figure 4C, Figure EV4A)**. Crucially, both wild type and S1121A separase became activated only after mimicking fertilization and not before, indicating that the injected separase was functional and activity was correctly controlled.

### Securin is not an essential inhibitor of separase in meiosis II

As shown before, securin is not required during the transition from meiosis I into meiosis II to keep separase under control, and *securin^−/−^* oocytes progress into metaphase II where they remain arrested to await fertilization, without showing significant precocious sister chromatid segregation during the arrest **(Figure 2D and E)**. To create conditions where separase is only inhibited by cyclin B1 and not securin, we used double knock-out *sep^−/−^ securin^−/−^* oocytes that were allowed to progress into metaphase II. In metaphase II we expressed wild type separase, which can still be inhibited by cyclin B1 (**Figure 5A**). When wild type separase was expressed in *sep^−/−^ securin^−/−^* metaphase II oocytes, the absence of securin in metaphase II did not cause chromosome or sister chromatid separation (**Figure 5B (top right), C, D (bottom), and E)**. This demonstrates that on the contrary to (Nabti et al., 2008), cyclin B1 on its own is sufficient to inhibit separase, and securin is not the major separase inhibitor in metaphase II. We asked whether the amount of exogenous separase was sufficient for cohesin cleavage in the metaphase II rescue experiment. Inducing anaphase II onset by strontium treatment resulted in the timely segregation of chromosomes, indicating that enough active separase was expressed **(Figure 5C, Figure EV4)**.

**Figure 5.**
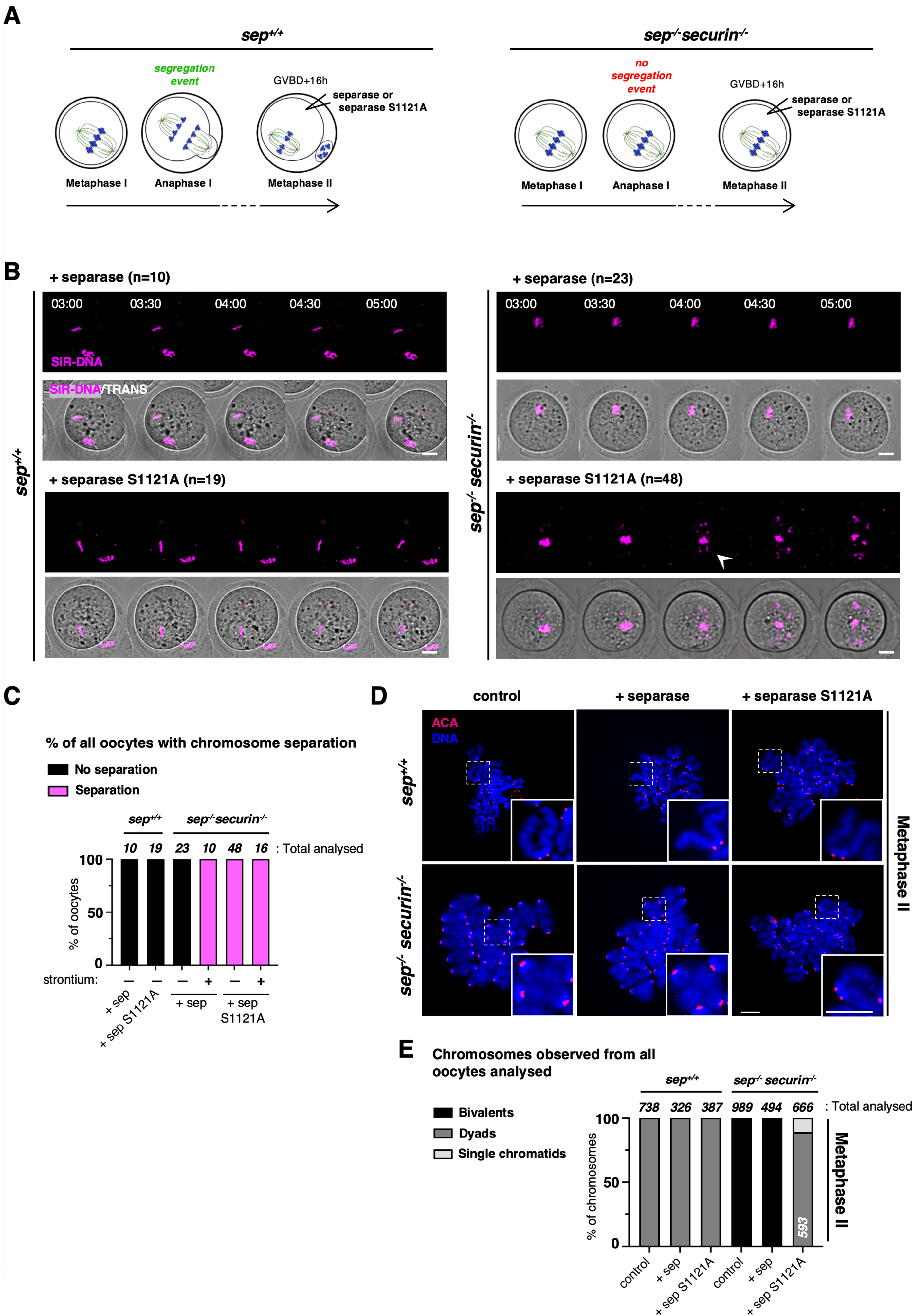
Loss of separase control by interfering with both cyclin B1 and securin in meiosis II. **A)** Illustrative scheme of the experimental setup: *sep^+/+^* (*separase^LoxP/LoxP^*) and *sep^−/−^securin^−/−^* oocytes were matured *in vitro* until metaphase II and injected with mRNAs encoding for separase or separase S1121A. Post injections, oocytes were subjected to live imaging to track either chromosome segregation (B), or chromosome spreads were performed to verify cohesin cleavage (D). **B)** Live imaging of *sep^+/+^* (left) and *sep^−/−^securin^−/−^* (right) oocytes expressing separase or separase S1121A where indicated, such as described in (A). For all conditions, oocytes were injected around 16h after GVBD (arrested in metaphase II) and were subjected to live imaging around 18h after GVBD. Time after start of the movie is shown in hours:minutes (separation of chromosomes is indicated with an arrow head). Prior to acquisition, oocytes were preincubated in culture media containing SiR-DNA to visualize chromosomes. Efficient expression of the separase constructs was verified by inducing anaphase II chemically with strontium. n is the number of oocytes analysed. Scale bar (white) represents 20 μm. (**Related to Figure EV4A and movie EV3**). **C)** Frequency of DNA separation observed in metaphase II-arrested oocytes, indicating separase activation, in the corresponding conditions of the selected time frames shown in (B). The total number of oocytes analysed for each condition, and of each category, is indicated. sep: separase. (**Related to Figure EV4A**) **D)** *sep^+/+^* (top) and *sep^−/−^securin^−/−^* (bottom) oocytes were injected around 16h after GVBD (arrested in metaphase II) with the indicated constructs, and were fixed for chromosome spreads around 22h after GVBD (Metaphase II). Centromeres/kinetochores were stained with ACA (red) and chromosomes with DAPI (blue). A representative spread and magnification of one chromosome (white dashed line squares, insert at the bottom right corner) are shown for each condition. Total number of chromosomes analysed is indicated in (E). Scale bars (white) represents 10 μm. **E)** Frequency of chromosome categories observed at metaphase II arrest (22h after GVBD), quantified from chromosome spreads shown in (D). Total number of chromosomes quantified for each condition, and of each category, is indicated. sep: separase.

### Meiosis II without separase inhibition

To determine whether centromeric cohesin is still protected during the prolonged fertilization arrest in metaphase II, we wanted to remove all separase inhibitors. As before in meiosis I, we asked whether loss of both cyclin B1 and securin inhibition of separase leads to constitutively active separase, independently of cell cycle progression into anaphase II. If not, a third inhibitor such as Sgo2-Mad2 may contribute to separase inhibition in meiosis II and keep separase inactive (Hellmuth et al., 2020). *sep^−/−^ securin^−/−^* oocytes in metaphase II were rescued with separase S1121A to obtain active separase, decoupled from cell cycle progression, in meiosis II. Indeed, such as in meiosis I, separase S1121A is not inhibited and cleaves cohesin even though oocytes have not been fertilized or activated with strontium **(Figure 5A, B (bottom right), C, D (bottom right), and E, EV4)**. But because *sep^−/−^ securin^−/−^* oocytes progressed until metaphase II without separase activity and thus, without kinetochore individualization, loss of inhibition of separase that was exogenously expressed in metaphase II led to mostly chromosome separation, and hardly any sister chromatids were observed. The resulting dyads had individualized kinetochores, such as previously observed when wild type separase was expressed from metaphase II onwards in *sep^−/−^* oocytes treated with strontium to undergo anaphase II **(Figure 5D and E)**. We found previously that sister kinetochore individualization has to take place in the previous division for centromeric cohesin cleavage, indicating that individualization and centromeric cohesin cleavage must be temporally separated (Gryaznova et al., 2021). Thus, our experimental setting allowed us to obtain constitutively active separase in metaphase II and confirms the importance of kinetochore individualization for deprotection, but did not allow us to address whether cohesin protection after kinetochore individualization is still present in metaphase II.

### Metaphase II without separase inhibition reveals absence of cohesin protection

Having determined how separase inhibition can be abolished in meiosis II allowed us to attempt answering the long-standing question of whether centromeric cohesin protection is still present throughout the extended metaphase II arrest in oocytes that harbor the usual dyads. We used *securin^−/−^* oocytes, which progress normally through meiosis I with the segregation of chromosomes and kinetochore individualization due to the activity of endogenous separase. We expressed S1121A separase in addition to endogenous separase in metaphase II *securin^−/−^* oocytes **(Figure 6A)**. Importantly, we found that expression of S1121A separase (which is inhibited neither by cyclin B1 nor securin in this context) led to immediate sister chromatid separation of nearly all dyads **(Figure 6B, C)**. This result indicates that centromeric cohesin protection is absent in metaphase II, but only if kinetochore individualization has previously taken place. Our result shows that even though Sgo2 and PP2A are present at metaphase II kinetochores and centromeres (Chambon et al., 2013, Gryaznova et al., 2021, Mengoli et al., 2021), they do not confer efficient centromeric cohesin protection.

**Figure 6.**
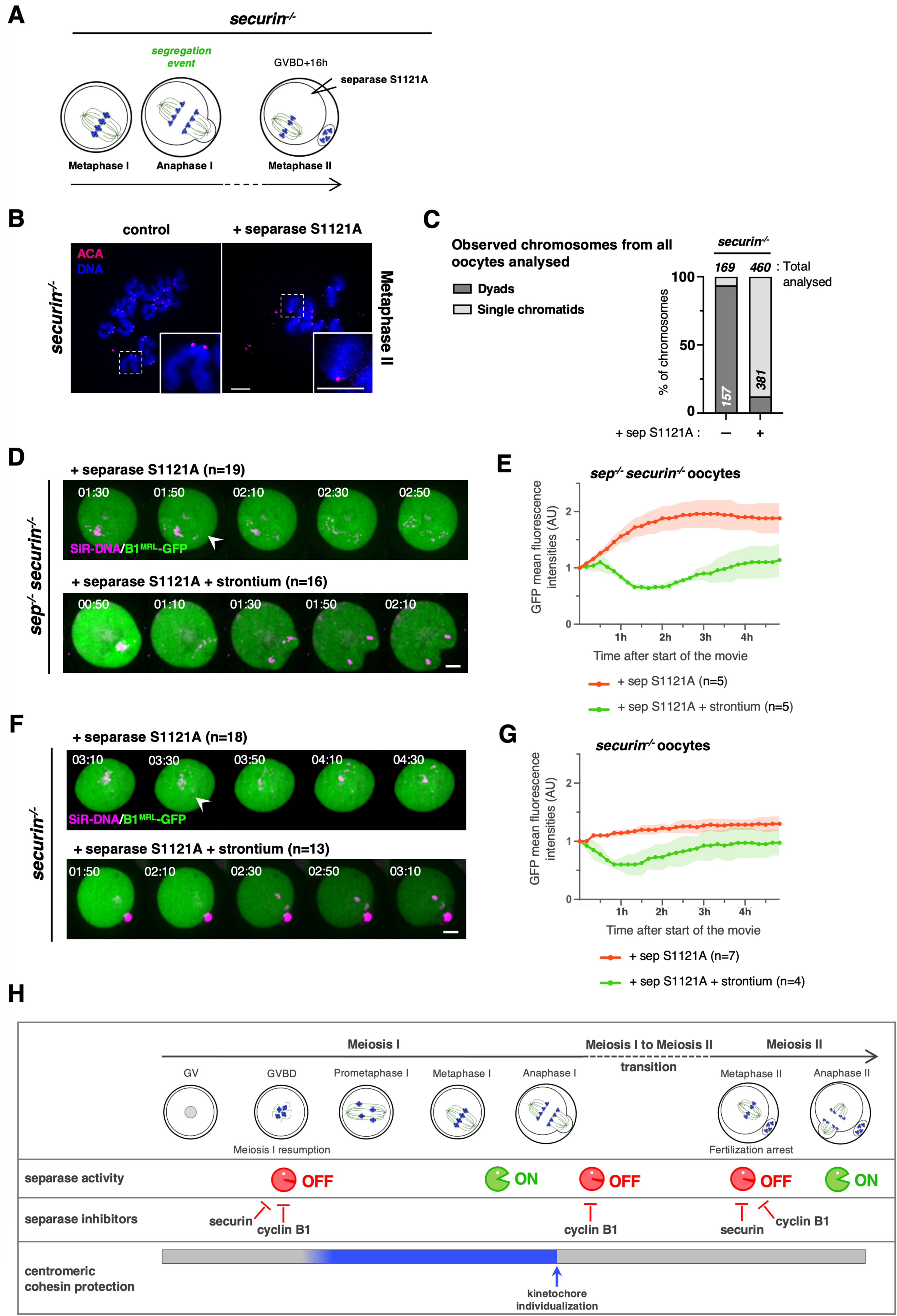
Loss of separase inhibition by cyclin B1 and securin in meiosis II reveals absence of efficient cohesin protection. **A)** Illustrative scheme of the experimental setup: *securin^−/−^* oocytes were matured *in vitro* until metaphase II and injected with mRNAs encoding for separase S1121A around 16h after GVBD, such as described in Fig 4A. **B)** *securin^−/−^* oocytes expressing S1121A separase, where indicated, were fixed for chromosome spreads around 22h after GVBD (Metaphase II). Chromosome spreads were stained for centromeres/kinetochores were stained with ACA (red) and chromosomes with DAPI (blue). A representative spread and magnification of one chromosome (white dashed line squares, insert at the bottom right corner) are shown for each condition. Total number of chromosomes analysed is indicated in (C). Scale bars (white) represents 10 μm. **C)** Frequency of chromosome categories observed at metaphase II arrest (22h after GVBD), quantified from chromosome spreads shown in (B). Total number of chromosomes quantified for each condition, and of each category, is indicated. **D)** Overlays of live imaging of *sep^−/−^securin^−/−^* double knock-out oocytes expressing separase S1121A and the GFP-cyclin B1 MRL mutant. Oocytes were injected with the constructs around 16h after GVBD, and activated with strontium, where indicated. Oocytes were then subjected to live imaging around 18h after GVBD, corresponding to 5 minutes after activation with strontium. Time after start of the movie is shown in hours:minutes (separation of chromosomes is indicated with an arrow head). (**Related to Figure EV5**). **E)** Quantification of GFP-cyclin B1 MRL in the corresponding conditions of the selected time frames shown in (D). Hours after start of the movie are indicated, error bars are ± SD, and n is number oocytes used for quantification. Shown, quantification of one representative replicate. **F)** Overlays of live imaging of *securin^−/−^* knock-out oocytes activated with strontium, where indicated, expressing separase S1121A and the GFP-cyclin B1 MRL mutant such as described in (D). Time after start of the movie is shown in hours:minutes (separation of chromosomes is indicated with an arrow head). (**Related to Figure EV5**). **G)** Quantification of GFP-cyclin B1 MRL in the corresponding conditions of the selected time frames shown in (F). Hours after start of the movie are indicated, error bars are ± SD, and n is number oocytes used for quantification. Shown, quantification of one representative replicate. **H)** Model of how separase activity and centromeric cohesin protection are coordinated during meiotic cell cycle progression in oocytes. See text for details.

### APC/C activation is not required for sister chromatid segregation in meiosis II

In budding yeast, deprotection of centromeric cohesin was shown to depend on APC/C activation, interfering with Sgo1-PP2A protective function (Jonak et al., 2017, Mengoli et al., 2021). We were wondering whether APC/C activation was equally required for sister chromatid segregation in metaphase II oocytes. In other words, was the cleavage of centromeric cohesin upon S1121A separase expression in *securin^−/−^* oocytes due to spontaneous anaphase II onset, or did it take place independently of cell cycle progression? In metaphase II-arrested oocytes, all kinetochores are attached, thus we could not use SAC satisfaction as a readout for APC/C activation, such as in meiosis I. To address this question, and without interfering with separase activity, we followed degradation of a cyclin B1 mutant carrying a fluorescent tag, which is devoid of kinase activity due to mutations in the MRAIL motif (required for correct binding to Cdk1 (Jeffrey et al., 1995, Schulman et al., 1998)). As expected, *securin^−/−^* and *sep^−/−^ securin^−/−^* oocytes that were activated with strontium showed degradation of cyclin B1-MRL-GFP. However, *securin^−/−^* and *sep^−/−^ securin^−/−^* oocytes injected with S1121A separase and not subjected to activation, showed separated sister chromatids or chromosomes, respectively, but did not degrade cyclin B1-MRL-GFP. **(Figure 6D-G)**. No proper anaphase movements, nor any attempts of second PB extrusion were observed, further showing that oocytes are still cell cycle arrested (**Figure EV5, Movie EV3**). Thus, we conclude that cleavage of centromeric cohesin and sister chromatid segregation in oocyte meiosis II only depends on kinetochore individualization and separase activation. Constitutive active separase can cleave centromeric cohesin without APC/C activation counteracting Sgo2-PP2A.

## Discussion

Separase cleaves the kleisin subunit of the cohesin complex, and its inappropriate activation leads to cohesin loss with weakened physical connections holding sister chromatids together. In the worst case, sister chromatids will separate precociously, leading to both mitotic and meiotic aneuploidies. Separase could thus be categorized as a “dangerous” enzyme requiring balanced control. This is also reflected by the fact that intracellular securin homeostasis is important for maintaining integrity of pituitary cell adenoma formation (Chesnokova et al., 2008). In mitosis, the time when separase could inappropriately cleave mitotic Scc1 and thus, weaken cohesion, may be rather limited, as before entry into mitosis, separase is localized to the cytoplasm and physically separated from cohesin. Until metaphase-to-anaphase transition, separase is kept under control by three independent mechanisms: 1) securin, which functions as a pseudo-substrate inhibitor and at the same time, is required for full separase activity through its chaperone activity; 2) cyclin B1-Cdk1, which phosphorylates separase and creates a pocket binding site to bind separase, resulting in inhibition of separase by cyclin B1, but equally, of cyclin B1-Cdk1’s kinase activity by separase; and 3) Sgo2-Mad2, proposed to bind separase and inhibit its cleavage activity in a pseudo-substrate dependent manner. Exit from the mitotic cell cycle with the reformation of a nuclear envelope and nuclear exclusion of separase protects cohesin from being cleaved until the subsequent mitosis (Wassmann, 2022, Yu et al., 2023).

### Separase inhibition until metaphase I

Our study was motivated by the quest to determine when, during oocyte meiosis I and II, centromeric cohesin is protected from cleavage by separase. For this, it was necessary to generate oocytes containing separase that is constitutively active by removing its inhibitors. We found that in oocytes, either securin or cyclin B1 is sufficient to maintain separase under control until metaphase I. At the transition from meiosis I into meiosis II, cyclin B1 plays a major role in separase inhibition, suggesting a hand-over from securin-dependent inhibition to cyclin B1, such as proposed in mitosis (Yu et al., 2023). This premise fits well with the fact that cyclin B1 degradation occurs in a delayed manner compared to securin, and that some cyclin B1 protein can still be detected at exit from meiosis I, even though Cdk1 kinase activity is undetectable (Karasu et al., 2019, Levasseur et al., 2019, Niault et al., 2007, Thomas et al., 2021). The complete loss of separase control shows that Sgo2-Mad2 is not essential in oocyte meiosis for separase control. Sgo2-Mad2 cannot substitute for loss of securin and cyclin B1 in this respect, and if Sgo2-Mad2 has a role, it is likely redundant with the two other inhibitors. Sgo2-Mad2 may become important once protection has been set up. In early meiosis I Sgo2 may not yet have been recruited to sufficient high levels yet-not only for protection, but also to inhibit separase. However, to address the role of Sgo2-Mad2 convincingly, it will be necessary to invalidate Sgo2 or Mad2 specifically in metaphase I and not before, and find ways to separate a potential role in separase inhibition from other essential functions both proteins play in oocyte meiosis, such as cohesin protection, chromosome alignment and SAC control.

### Protection of centromeric cohesin is not yet set up when oocytes resume meiosis I

As all cohesin is cleaved in meiosis I when both securin and cyclin B1 as inhibitors are absent when oocytes resume meiosis, this implies that pericentromeric cohesin protection is set up rather late during meiosis I (**Figure 6H**). This correlates with our results showing that Sgo2 and PP2A are reduced at the centromere region at meiotic resumption and in early prometaphase I. Of note, oocytes devoid of spindle assembly checkpoint components are strongly accelerated in meiosis I progression and undergo anaphase I around 4-5 hours after GVBD. But even if these oocytes missegregate chromosomes, this does not lead to loss of protection. (One exception is the checkpoint kinase Mps1, which also contributes to Sgo2 localization for protection.) (El Yakoubi et al., 2017, McGuinness et al., 2009, Touati et al., 2015). Hence, protection of cohesin is functional at least 4-5 hours after meiosis resumption. Thus, we identified a critical moment at the start of oocyte maturation that may contribute to missegregations and thus the generation of aneuploidies, when tight separase control is perturbed.

### Transition from meiosis I into meiosis II

Another critical moment in oocyte maturation is the transition from meiosis I into meiosis II, because no nuclear envelope -to separate separase from Rec8-is reformed during this cell cycle transition. Additionally, separase inhibition depends only on cyclin B1 and not securin. The fact that Sgo2-PP2A become strongly reduced at the centromeric region as oocytes progress from meiosis I into meiosis II previously prompted us to ask whether an additional mechanism prevents separase from accessing pericentromeric cohesin. Indeed, we found that sister kinetochore individualization is required for deprotection and cleavage of pericentromeric cohesin (Gryaznova et al., 2021). However, this individualization takes place in anaphase I, and thus well before reaccumulation of securin and cyclin B1. It remains enigmatic how centromeric cohesin cleavage is avoided once sister kinetochore individualization has taken place, and cyclin B1 and securin are still absent to re-inhibit separase. As an additional control to avoid that separase cleaves pericentromeric cohesin once cyclin B1 is degraded, the kinase phosphorylating Rec8 for cleavage in meiosis II may be localized away from Rec8. Indeed, Aurora B/C, the kinase(s) promoting Rec8 phosphorylation in meiosis I, are part of the chromosomal passenger complex and thus localized to the spindle midzone (Nguyen & Schindler, 2017, Nikalayevich et al., 2022).

It has been proposed that cyclin B1-Cdk1 activity only partially recedes between meiosis I and meiosis II, allowing certain substrates to remain phosphorylated during the transition and thus, preventing exit into interphase. It remains to be seen whether residual cyclin B1-Cdk1 activity must be retained to maintain separase inactive. In mitotic cells separase autocleavage, which takes place when separase is liberated from securin binding, removes a binding site for PP2A. As a result, cyclin B1 phosphorylation of separase is not counteracted by PP2A binding to separase, thus promoting inhibition of separase by cyclin B1-Cdk1 binding, as long as cyclin B1 protein is still present (Hertz et al., 2016, Holland et al., 2007, Shindo et al., 2022, Yu et al., 2023). We suggest that similar to securin (Thomas et al., 2021), cyclin B1 protein bound to separase (devoid of its PP2A binding site) is not degraded efficiently in oocytes. Cyclin B1 may thus inhibit separase during the short time window when APC/C is active and cyclin B1-Cdk1 associated activity absent, before re-accumulation of cyclin B1 in prometaphase of meiosis II. This would explain why some cyclin B1 protein is still present in oocytes exiting meiosis I (Karasu et al., 2019). In a scenario where separase would remain phosphorylated due to autocleavage of the PP2A binding site, cyclin B1 might remain associated with separase long enough to inhibit separase until entry into meiosis II. As cyclin B1-Cdk1 and separase mutually inhibit each other (Gorr et al., 2005), no Cdk1-associated kinase activity is detected (Karasu et al., 2019). This hypothesis is consistant with our result that precocious sister chromatid segregation is observed in oocytes that progress into meiosis II without cyclin B1-dependent inhibition of separase (**Figure 1G and H**). However, more experiments are needed to confirm or invalidate this hypothesis.

### Cohesin is already deprotected during the oocyte metaphase II arrest

In budding yeast, artificial activation of separase in metaphase II-arrested cells (due to conditional inhibition of the APC/C activator Cdc20) does not lead to centromeric cohesin removal. Additional removal of Sgo1 was necessary to deprotect Rec8 and induce sister chromatid segregation (Mengoli et al., 2021). Thus, active separase is not sufficient in budding yeast metaphase II to cleave centromeric Rec8, presumably because protection mediated by Sgo1 is still present and also needs to be removed. This is in contrast with our findings here in mouse oocytes, where elimination of both cyclin B1- and securin-dependent inhibition of separase led to immediate separation of sister chromatids in metaphase II. We expected that APC/C activation would still be required to remove cohesin protection at this time such as in budding yeast. However, we found here that cleavage of centromeric cohesin took place without APC/C activation, or visible anaphase movements. Hence, Sgo2 and PP2A localized to the centromere region were apparently unable to protect centromeric cohesin in the presence of active separase at this stage of the cell cycle. Crucially, Sgo2 and PP2A are also involved in other functions at the kinetochore/centromere in oocytes (Rattani et al., 2013). Therefore, their sole presence does not indicate that protection is indeed functional, at least in meiosis II. The only condition for cleavage of pericentromeric cohesin in meiosis II in oocytes was previous bivalent segregation. This finding is in accordance with our model that sister kinetochore individualization occurring at meiosis I exit is essential for separase to access pericentromeric cohesin, but only in the following division, meiosis II (Gryaznova et al., 2021). We speculate that cohesin protection is still present in budding yeast metaphase II, as in this model system, only Pds1 (securin) functions as a separase inhibitor (Kamenz & Hauf, 2017, Mengoli et al., 2021). Yeast meiosis may thus additionally rely on protection of centromeric cohesin to avoid precocious sister chromatid segregation. In mouse oocytes, inhibition of separase by securin and cyclin B1-Cdk1, and distinct localization of the kinase phosphorylating Rec8 likely substitutes for protection in meiosis II.

In conclusion, our results show that either of the main known inhibitors of vertebrate separase - namely securin or cyclin B1-Cdk1-is sufficient to maintain separase under control, except at the transition from meiosis I into meiosis II, which depends mainly on cyclin B1-Cdk1. Furthermore, we found that centromeric cohesin protection is in place only during a relatively short time window during meiosis I (**Figure 6H**). Tight control of separase is the main factor preventing precocious centromeric cohesin removal as oocytes progress from meiosis I into meiosis II. This study thus elucidates critical moments during oocyte maturation when aneuploidies may arise in vertebrates. It remains intriguing that oocyte meiosis does not depend on additional mechanisms to avoid untimely cohesin removal throughout meiotic maturation and metaphase II arrest.

### Reagents and Tools Table

| Reagent/Resource | Reference or Source | Identifier or Catalog Number |
| --- | --- | --- |
| **Experimental Models** |  |  |
| C57Bl6/J (*M. musculus*) | Janvier Labs | C57BL/6-Jrj |
| Swiss (*M. musculus*) | Janvier Labs | RjOrl:SWISS (CD-1) |
| **Recombinant DNA** |  |  |
| pRN3-H2B-mCherry-hScc1(107-268)-YFP | *this study* | Nikalayevich *et al* (2022) |
| pRN3-mseparase | Kudo *et al* (2006) |  |
| pRN3-mseparase S1121A | Touati *et al* (2012) |  |
| pRN3-human Cyclin B1(MRL)-mGFP | *this study* |  |
| **Antibodies** |  |  |
| Human anti-centromere protein (ACA) | Clinisciences | 15-234 |
| Rabbit anti-Sgo2 | *this study* |  |
| Mouse anti-PP2AC-1D6-Alexa Fluor 488 | Sigma-Aldrich | 05-421-AF488 |
| Rabbit anti-Mad2 | (Wassmann, 2022) *et al* (1998) |  |
| Donkey anti-human CY3 | Jackson Immuno Research | 709-166-149 |
| Donkey anti-human Alexa Fluor 647 | Jackson Immuno Research | 709-606-149 |
| Donkey anti-rabbit CY3 | Jackson Immuno Research | 711-166-152 |
| Donkey anti-rabbit Alexa Fluor 488 | Jackson Immuno Research | 711-546-152 |
| Donkey anti-mouse Alexa Fluor 488 | Jackson Immuno Research | 715-546-150 |
| **Chemicals, Enzymes and other reagents** |  |  |
| dibutylryl cyclic AMP | Sigma-Aldrich | D0260 |
| Embryo culture oil | FUJIFILM Irvine Scientific | 9305 |
| Strontium chloride hexahydrate | Sigma-Aldrich | 204463 |
| Paraformaldehyde | Sigma-Aldrich | 441244 |
| Triton X-100 | Sigma-Aldrich | T8787 |
| Dithiothreitol (DTT) | Sigma-Aldrich | D9779 |
| Bovine Serum Albumin (BSA) | Sigma-Aldrich | A4378 |
| Vectashield DAPI | EUROBIO Scientific | H-1200 |
| SiR-DNA | Spirochrome | SC007 |
| Nocodazole | Sigma-Aldrich | M1404 |
| **Software** |  |  |
| Metamorph 7.10.3 | Metamorph | https://support.moleculardevices.com/s/article/MetaMorph-Software-installation-files |
| Fiji 5.1.0 | Fiji | https://imagej.net/software/fiji/ |
| PRISM 10 | GraphPad | https://www.graphpad.com/scientific-software/prism/ |
| **Other** |  |  |
| mMessage mMachine T3 kit | Invitrogen | AM1348 |
| RNeasy Mini Kit 5 | Qiagen | 74104 |

## Material and Methods

### Animals

Mice were kept in an enriched environment with *ad libitum* food and water access in the conventional animal facility of UMR7622 (authorization C 75-05-13) and UMR7592 (authorization C75-13-17), according to current French guidelines. Housing was done under a 12-hour light / 12-hour dark cycle according to the Federation of European Laboratory Science Associations (FELASA). All experiments were subject to ethical review and approved by the French Ministry of Higher Education and Research (authorization n° B-75-0513), in accordance with national guidelines and application of the “3 Rs” rule (Licence 5330). The number of mice used was kept as low as possible but high enough to obtain statistically significant results, in agreement with the standards used in the field. Females for experiments were either house-bred in the animal facility of UMR7592 (genetic knock-out and littermate control mice), or purchased at 7 weeks of age (C57BL/6-Jrj and Swiss, Janvier Labs France) and kept until full sexual maturity. For all experiments, mice were euthanized by cervical dislocation between 9 and 16 weeks of age, by trained personal. Mice have not been involved in any procedure except genotyping prior euthanasia.

### Generation of mouse strains, husbandry, and genotyping

*separase^LoxP/LoxP^, separase^LoxP/LoxP^ Zp3 Cre^+^* (Kudo et al., 2006), *Pttg^−/−^* (Wang et al., 2001), and *separase^LoxP/LoxP^ Zp3 Cre^+^ Pttg^−/−^* mice were house-bred in the animal facilities of UMR7622 (*separase^LoxP/LoxP^ Zp3 Cre^+^* and *Pttg^−/−^* strain) and UMR7592 (mice of all three strains used for experiments in this study). Primer sequences used for genotyping are available upon request. Littermates were used as controls in experiments. Replicate experiments were performed with mice from different litters.

### Mouse oocyte harvesting and *in vitro* culture

Fully grown prophase I-arrested oocytes were harvested from dissected ovaries. Oocytes were retrieved by mouth pipetting follicles through narrow hand-pulled glass pasteur pipets. Selected fully grown prophase I-arrested oocytes were cultured at 38°C in drops of self-made M2 media supplemented with 100 µg/mL dbcAMP (dibutyryl cyclic AMP, Sigma-Aldrich, D0260) and covered with mineral oil (suitable for embryo culture, FUJIFILM Irvine Scientific, 9305). To induce resumption of meiosis I from prophase I, and meiotic maturation, oocytes were washed three times in abundant drops of M2 media. Oocytes were synchronized visually at GVBD (Germinal vesicle breakdown), and only oocytes undergoing GVBD within 90 minutes after release were used (El Jailani et al., 2024). For meiosis II experiments, oocytes were always harvested in M2 media and then cultured overnight in M16 media (homemade).

To study Mad2 recruitment to kinetochores as a read-out of APC/C activity, oocytes were treated with the spindle-depolymerizing drug nocodazole (Sigma-Aldrich, M1404). Oocytes were cultured in M2 media supplemented with nocodazole, from GVBD until chromosome spreads were performed, at a final concentration of 400nM.

For activation experiments in meiosis II, oocytes were cultured in M2 media after overnight incubation in M16. Strontium chloride (SrCl2, Sigma-Aldrich, 204463) was used to release oocytes from metaphase II arrest. At 16,5 hours after GVBD, oocytes of the indicated genotypes were incubated in M2 medium without CaCl2 (activation medium, homemade) for 1 hour, and transferred into activation medium supplemented with 100mM SrCl2 (Bouftas et al., 2022). After washing the oocytes in 6 droplets of activation medium, they were observed by live imaging microscopy to visualize release from CSF arrest.

### In vitro transcription and oocyte microinjection

Plasmids used are described in the Reagents and Tools Table. The cleavage sensor H2B-mCherry-Scc1-YFP construct (Nikalayevich et al., 2018) was cloned into a pRN3-T3 plasmid. Plasmids to express separase have been described (Kudo et al., 2006, Touati et al., 2012). The human cyclin B1-GFP plasmid (described previously in (Herbert et al., 2003)) was used as a template to generate a kinase-dead hydrophobic patch mutant (MRL mutant), using the Q5 site directed mutagenesis kit to mutate the MRAIL motif to AAAIA.

Primer sequences used for PCR cloning are available upon request. Plasmids were digested and linearized with the restriction enzyme Sfi1. Corresponding mRNAs were obtained by performing *in vitro* transcription using the mMessage mMachine T3 Kit (Invitrogen, AM1348). After 3h of transcription, mRNAs were purified by using the RNeasy Mini Kit 5 (Qiagen, 74104).

Purified mRNAs were injected into GV- or metaphase II-arrested oocytes. GV- and metaphase II-arrested oocytes were left for 1 hour to allow expression of protein before release from prophase I-arrest (GV-arrested oocytes) or start of the movie (with or without activation with strontium, metaphase II-arrested oocytes). Microinjection pipettes were made by pulling capillaries (Harvard Apparatus, 30-0038) with a magnetic puller (Next Generation Micropipette Puller P-1000, Sutter Instrument). Oocytes were manipulated on an inverted Nikon Eclipse Ti microscope equipped with Eppendorf micromanipulators, with a holding pipette (Eppendorf, VacuTip I EP5195000036-25, and Vitrolife, 15331), and injections were done using a FemtoJet Microinjection pump (Eppendorf) with continuous flow settings.

### Chromosome spreads

Tyrode’s acid treatment (El Jailani et al., 2024) was used to dissolve the *zona pellucida* (ZP) of oocytes. ZP-free oocytes were fixed in hypotonic solution containing 1% paraformaldehyde (Sigma-Aldrich 441244), 0,15% Triton X-100 (Sigma-Aldrich, T8787) and 3mM DTT (Sigma-Aldrich, D9779-1G) diluted in H20. Chromosome spreads were performed as described in (Bouftas et al., 2022), at the indicated time points, using microscopic 10-well slide (Fischer Scientific, 10588681).

### Immunostaining

To stain chromosome spreads, a first step of saturation was done with PBS supplemented with 3% BSA (Sigma-Aldrich, A4378) final concentration. Spreads were incubated at room temperature for 2 hours with primary antibodies, washed with PBS and incubated 1h with secondary antibodies. Centromeres were stained with human ACA primary antibody (anti-centromere protein antibody, Clinisciences, 15-234, 1:100). To stain for proteins involved in centromeric cohesion protection, we used rabbit polyclonal anti-Sgo2 generated against the last 20 aa in the C-terminus (self-made, 1:100), and mouse monoclonal anti-PP2A C-subunit clone 1D6 conjugated with Alexa Fluor 488 (Sigma-Aldrich, 05-421-AF488, 1:100). To stain Mad2, we used rabbit polyclonal anti-Mad2 (self-made, 1:150). The following secondary antibodies were used at 1:200: donkey anti-human Cy3 secondary antibody (Jackson Immuno Research, 709-166-149), donkey anti-human Alexa Fluor 647 (Jackson Immuno Research, 709-606-149), donkey anti-rabbit Cy3 (Jackson Immuno Research, 711-166-152), donkey anti-rabbit Alexa Fluor (Jackson Immuno Research, 711-546-152), donkey anti-mouse Alexa Fluor 488 (Jackson Immuno Research, 715-546-150). Slides were mounted with Vectashield mounting medium supplemented with DAPI (EUROBIO Scientific, H-1200). Acquisitions of stained chromosome spreads were made with either a Zeiss Axiovert 200M inverted microscope or a Nikon Eclipse Ti2-E inverted microscope, both coupled with an EMCD camera (Evolve 512, Photometrics), a Yokogawa CSU-X1 spinning disk, and equipped with appropriate lasers and filter-sets for three-color imaging, using a 100x/1.4 NA oil Dic Plan-Apochromat Zeiss objective or, 100x/1.45 oil Plan-Apochromat Nikon objective. 6 z-sections with 0,4mm interval were taken. Images were acquired by Metamorph software (Metamorph 7.10.3) and mounted with Fiji software (ImageJ2 2.9.0). No manipulations were made other than brightness and contrast adjustments that were applied with the same settings to images being compared.

### Live imaging

Live imaging of the cleavage sensor in mouse oocytes was performed under temperature-controlled conditions with a motorised inverted Nikon Eclipse Ti2-E microscope equipped with an EMCD camera (Evolve 512, Photometrics) or a high sensitivity sCMOS camera (Photometrics), and a PrecisExite High Power LED Fluorescence, using a 20x/0.75 NA oil Plan-Apochromat Nikon objective. Chambers with oocytes microinjected with mRNAs where indicated, were prepared for imaging as described in Nikalayevich et al (2018). Time lapse was set up for 16 hours in meiosis I, with time points taken every 20 mins. 15-20 z-sections with 3µm distance were acquired in mCherry and YFP channels for the cleavage sensor, and 1-z section for the TRANS image. For live imaging of mouse oocytes in meiosis II, a motorised inverted Nikon Eclipse Ti2-E microscope equipped with a Yokogawa CSU-X1 spinning disk, coupled with an EMCD camera (Evolve 512, Photometrics), with appropriate lasers and filter-sets for three-color imaging and a 20x/0.75 NA oil Plan-Apochromat Nikon objective was used. Time lapse was set up for 6h with 10 mins intervals. Prior acquisition, metaphase II oocytes were injected with mRNAs as indicated, pre-incubated in M2 media supplemented with 1mM SiR-DNA (far-red DNA labelling probe, Spirochrome, SC007) for 1 hour. 15-20 z-sections with 3µm distance were acquired in the 640nm channel for SiR-DNA, and 1-z section for the TRANS image. All time lapse acquisitions were piloted by Metamorph software (Metamorph 7.10.3). Stills of movies were mounted with Fiji software (ImageJ2 2.9.0) and contrast/brightness were adjusted equally to all conditions being compared.

### Quantification and statistical analysis

Quantifications were made using Fiji software. To quantify Sgo2, PP2A and Mad2 total signal on chromosome spreads, levels of fluorescence was measured on sum-projected images. A square of 10 x10 pixels was drawn on centromeres as depicted on the scheme in **Figure 3G** and **Figure 3I**. The signals measured (Ifluo) were corrected to background (bg) and normalized to ACA signals as follows: Ifluo(normalized) = [Ifluo - Ifluobg] / [IfluoACA - IfluoACA(bg)]. To quantify the mutant cyclin B1(MRL)-GFP degradation, mean levels of fluorescence intensities were measured on sum-projected time lapse images. A square of 100×100 pixels was drawn in the oocyte. Mean levels of fluorescence were measured for each time point of the time lapse movie and were corrected to background and normalized to the value of the first timepoint (t0). All data plots and statistical analysis were obtained with PRISM 10 software. To monitor separase activation timing, anaphase I onset was used as our read out. Error bars with standard deviation (SD), and mean bars are represented. Statistical analysis was performed using nonparametric Mann-Whitney U-test (ns: not significant, ** P<0.01, **** P< 0.0001). All experiments were done at least three times independently. Shown are results from all replicates performed or, of one representative replicate, where indicated.

## Acknowledgements

We thank lab members for discussions and useful suggestions, and Warif El Yakoubi for preliminary experiments. We thank Alberto Pendas (University of Salamanca) for providing *Pttg^−/−^* sperm straws, and Nicolas Minc (Institut Jacques Monod) for access to a Sutter Instruments Micropipette puller. We are grateful to Isabelle Le Parco and members of the animal house “Animalerie Buffon”, as well as the informatics and administrative support services of the Institut Jacques Monod. S.E.J. received a 4th year PhD fellowship from the Fondation pour la Recherche Médicale (Fin de thèse, FDT202404018087). Financial support for this work was obtained by the Fondation pour la Recherche Médicale (Equipe FRM DEQ 202103012574), and the Agence Nationale de la Recherche (ANR-23-CE13-0015-01) to K.W.

## Author contributions

**Safia El Jailani**: Data curation; Methodology; Formal analysis; Investigation; Visualization; Writing-original draft. **Damien Cladière**: Data curation; Formal analysis. Resources. **Elvira Nikalayevich**: Data curation; Formal analysis. **Sandra A. Touati**: Data curation; Formal analysis; Investigation. **Vera Chesnokova**: Resources. **Shlomo Melmed**: Resources. **Eulalie Buffin**: Supervision; Investigation. **Katja Wassmann**: Conceptualization; Supervision; Investigation; Funding acquisition; Writing-original draft.

## aExpanded View Figures

**Figure EV1 (related to Figure 1B).**
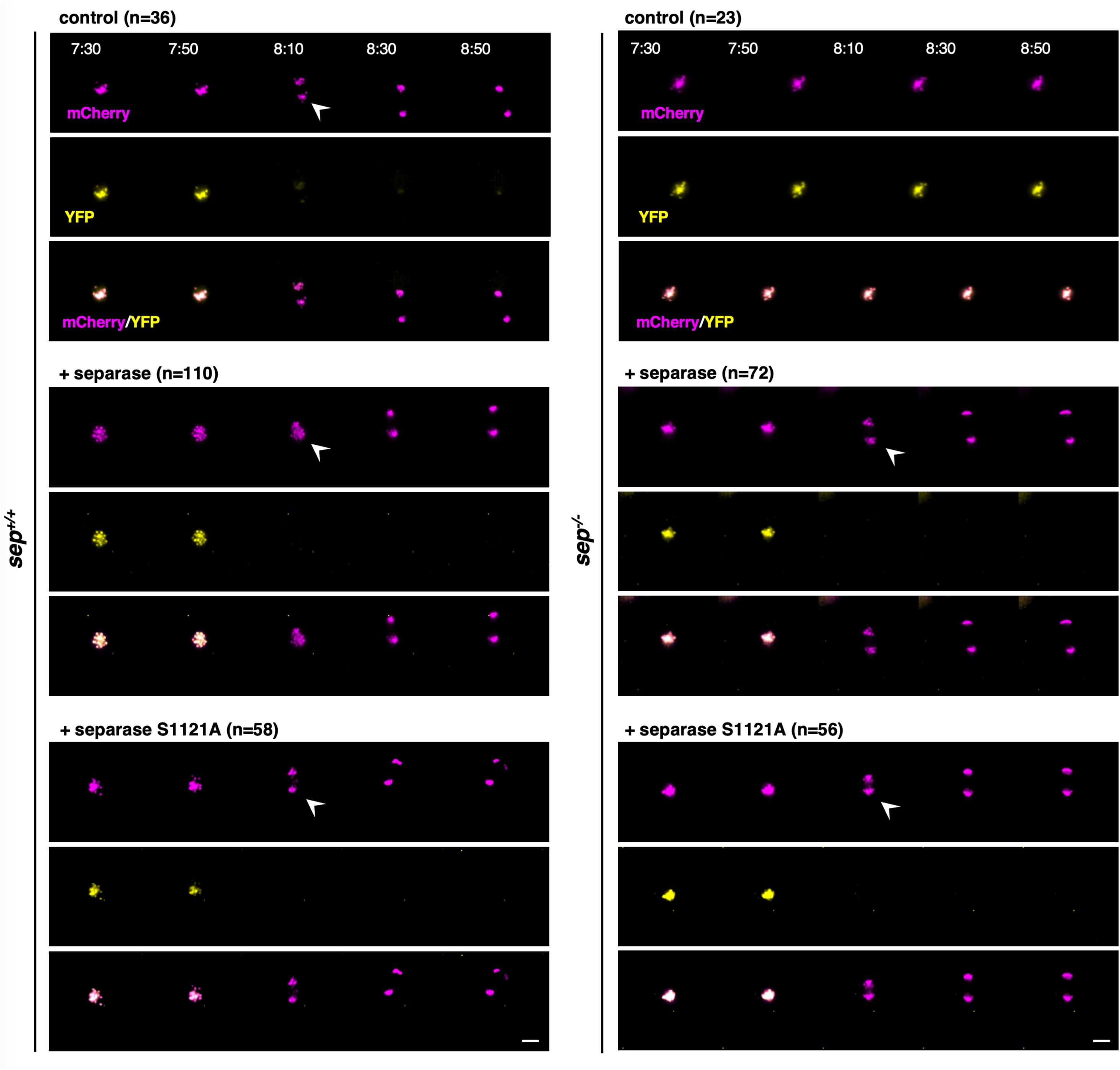
Cyclin B1 -dependent inhibition of separase is not essential before meiosis I exit. YFP and mCherry channels of the selected time frames overlays shown in Figure 1B. Time after GVBD is shown in hours:minutes (anaphase I onset is indicated with an arrow head). n is the number of oocytes analysed. Scale bar (white) represents 20 μm.

**Figure EV2 (related to Figure 2A).**
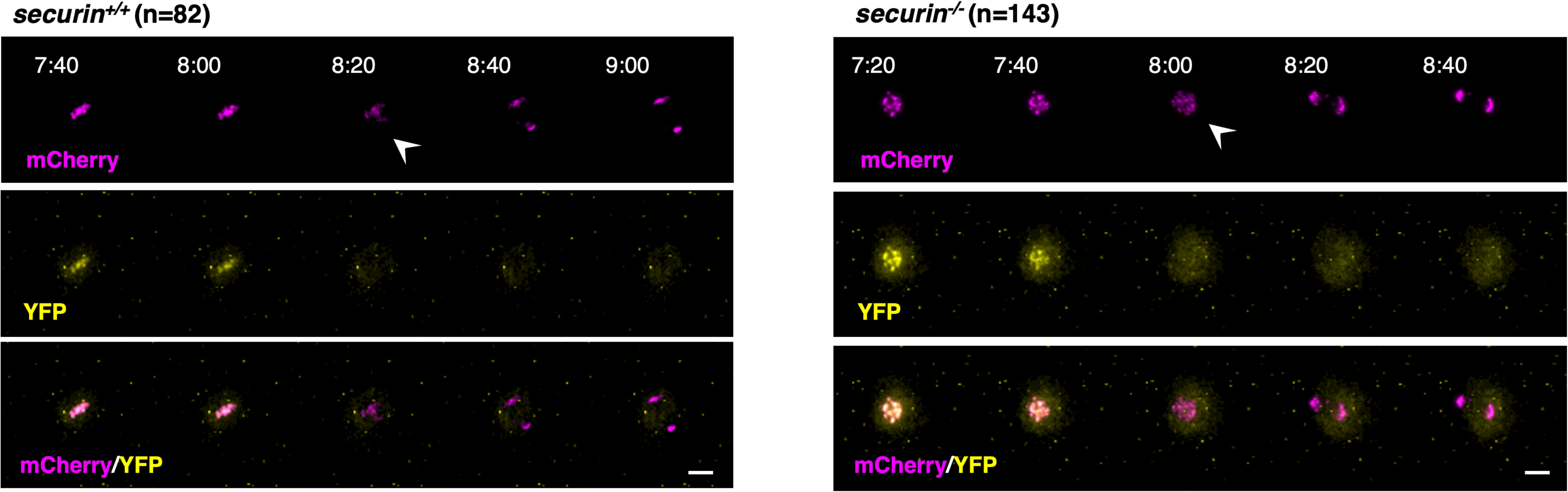
Securin is not essential for separase inhibition in meiosis. **I** YFP and mCherry channels of the selected time frames overlays shown in Figure 2A. Time after GVBD is shown in hours:minutes (anaphase I onset is indicated with an arrow head). n is the number of oocytes analysed. Scale bar (white) represents 20 μm.

**Figure EV3 (related to Figure 3A, movie EV2 and Appendix Figure S1).**
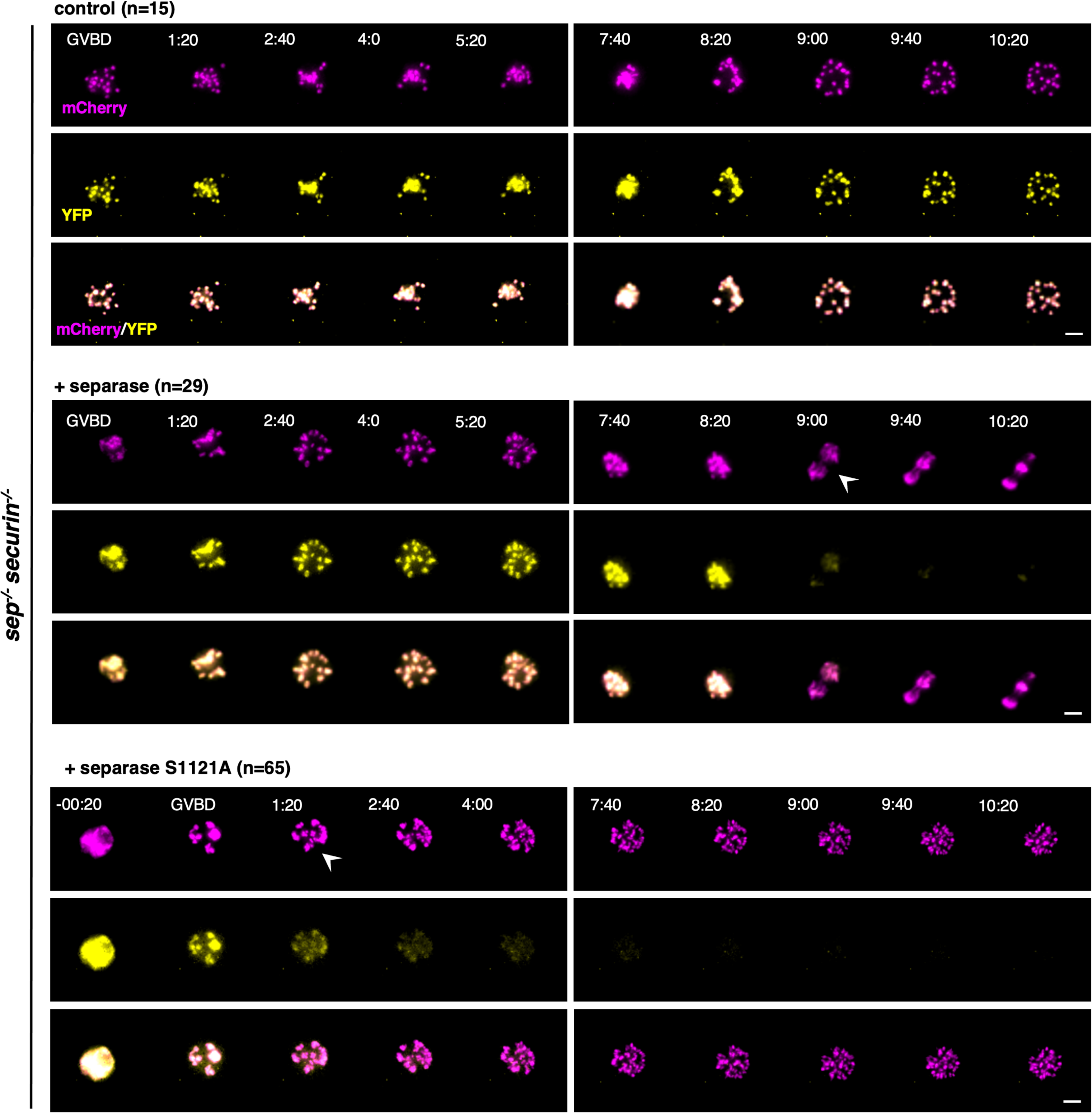
Complete loss of separase control reveals absence of cohesin protection in early prometaphase. **I** YFP and mCherry channels of selected time frames overlays shown in Figure 3A. Time after GVBD is shown in hours:minutes (cleavage onset is indicated with an arrow head). n is the number of oocytes analysed. Scale bar (white) represents 20 μm. Complete movies are shown in **Appendix Figure S1**.

**Figure EV4 (related to Figure 4B and 5B).**
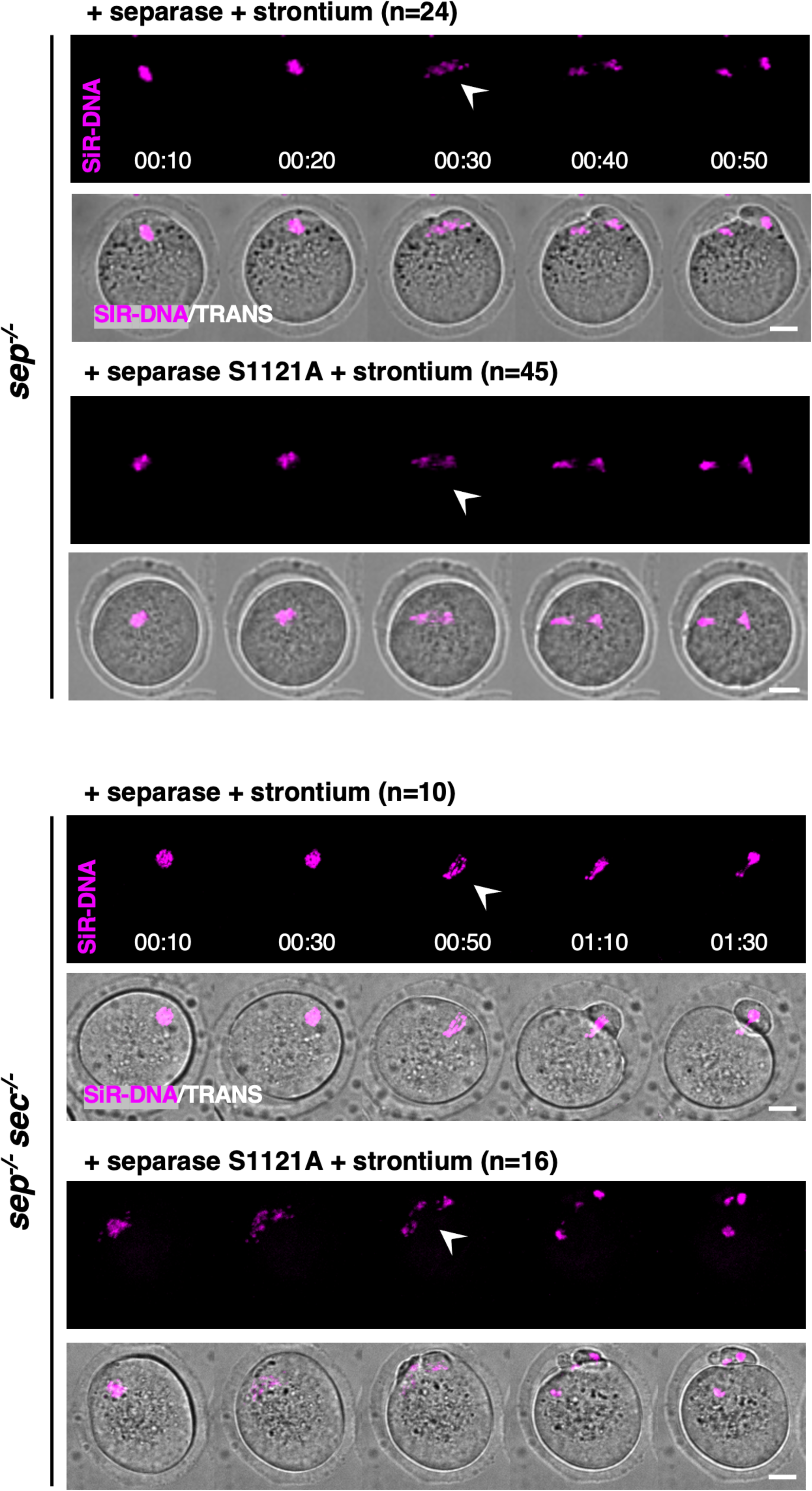
When rescued in metaphase II, *sep^−/−^* and *sep^−/−^securin^−/−^* oocytes can be activated to undergo meiosis II and separate chromosomes. Live imaging movies of *sep^+/+^* (top) and *sep^−/−^ securin^−/−^* (bottom) oocytes in Figure 4B **and 5B**. Where indicated, oocytes were chemically induced with strontium to verify sufficient expression of wild type separase or separase S1121A. For all conditions, oocytes were injected with the constructs around 16h after GVBD, and were subjected to live imaging around 18h after GVBD, corresponding to 5 minutes after oocytes were activated with strontium. Time after start of the movie is shown in hours:minutes (anaphase II onset is indicated with an arrow head). n is the number of oocytes analysed. Scale bar (white) represents 20 μm.

**Figure EV5 (related to Figure 6D and 6F).**
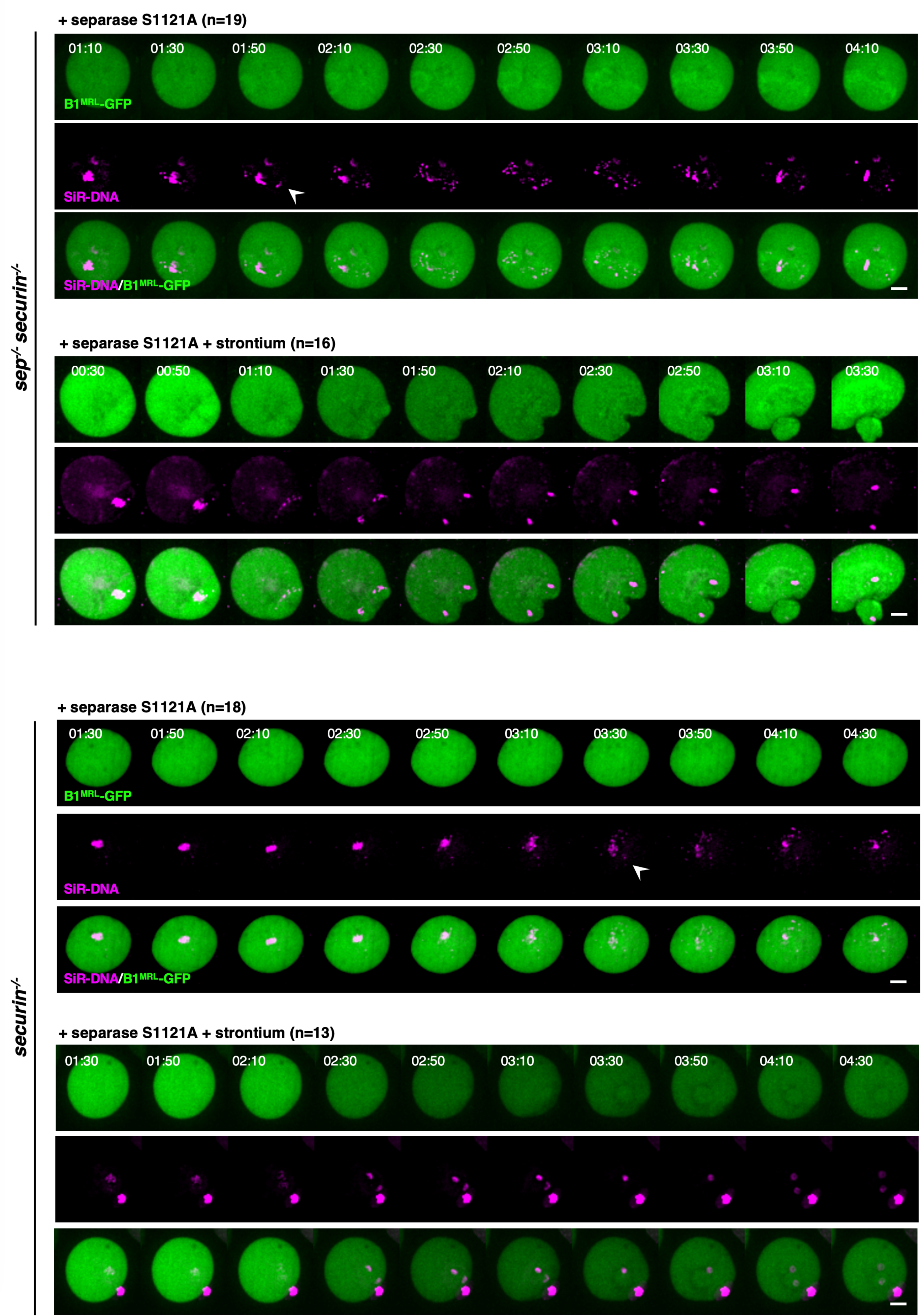
Separase out of control activation is independent of APC/C activation in meiosis II. Overlays of the GFP and Far-Red channels of the selected time frames shown in **Figure 6D and 6F**. *sep^−/−^ securin^−/−^* (top) and *securin^−/−^* (bottom) oocytes were injected with separase S1121A and the GFP-cyclin B1 MRL mutant around 16h after GVBD, and were subjected to live imaging around 18h after GVBD. Where indicated, oocytes were activated with strontium around 5 minutes before start of live imaging. Time after start of the movie is shown in hours:minutes (separation of chromosomes is indicated with an arrow head). n is the number of oocytes analysed. Scale bar (white) represents 20 μm.

## Expanded View Movies

**Movie EV1 (related to Figure 1B).**

Overlay of the YFP and mCherry channels of time lapse microscopy acquisitions of wild type (*sep^+/+^*) mouse oocytes expressing the cleavage sensor. Time after GVBD is shown in hours:minutes, and timepoints were taken every 20 mins, shown is the entire movie. Cleavage of the sensor is visible by the disappearance of the YFP signal from the chromosomes, whereas the mCherry signal remains localized to chromosomes. Scale bar (white) represents 20 μm.

**Movie EV2 (related to Figure 3A).**

Overlays of the YFP and mCherry channels of selected time frames shown in **Figure 3A**. *sep^−/−^ securin^−/−^* oocytes express the cleavage sensor and have been co-injected with mRNAs encoding for separase or separase S1121A, where indicated. Time after GVBD is shown in hours:minutes, and timepoints were taken every 20 mins, shown is the entire movie. Scale bar (white) represents 20 μm.

**Movie EV3 (related to Figure 5B).**

Time lapse acquisitions of the selected time frames shown in **Figure 5B** (bottom right). *sep^−/−^ securin^−/−^* oocytes were injected with separase S1121A around 16h after GVBD, and were subjected to live imaging around 18h after GVBD. Prior to acquisition, oocytes were preincubated in culture media containing SiR-DNA to visualize chromosomes. Time after start of the movie is shown in hours:minutes, shown is the entire movie. bar (white) represents 20 μm.

## Appendix

**Appendix Figure S1 (related to Figure 3A and Figure EV3).**
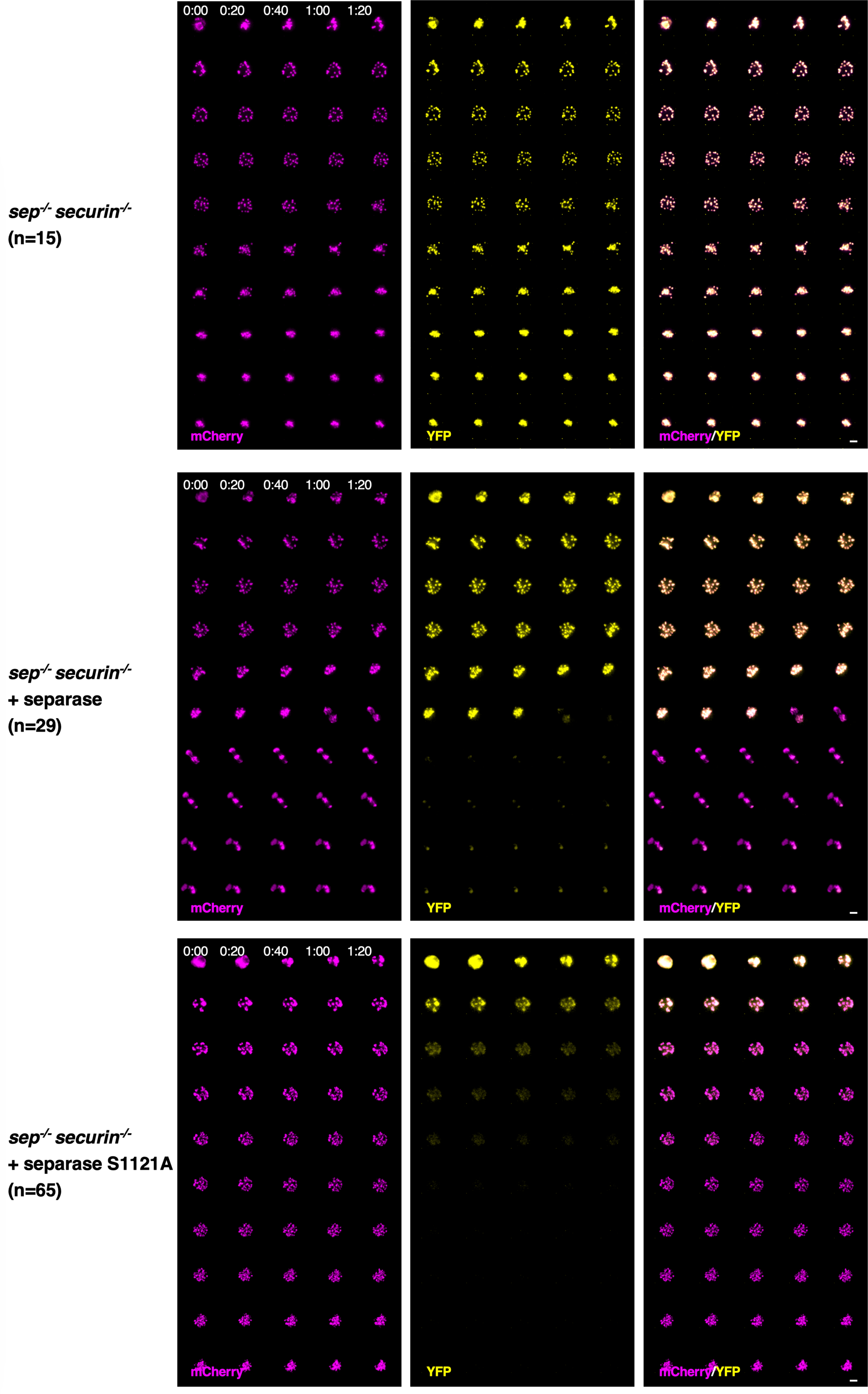
Complete loss of separase control reveals absence of cohesin protection in early prometaphase I. YFP and mCherry channels of selected time frames overlays shown in Figure 3A. Timepoints shown in hours (time after movie starts) and were taken every 20 mins, shown is the entire movie. n is the number of oocytes analysed. Scale bar (white) represents 20 μm.

